# Lysosomal Aspartylglucosaminidase Couples Glycoprotein Catabolism to Cell-Surface GlycoRNA Production

**DOI:** 10.64898/2026.01.13.699189

**Authors:** Wei Cao, Zehui Yang, Li Yi, Yulong Miao, Shuyang Zhao, Cheng Yu, Xiaoman Zhou, Ting Li, Yao Zhou, Tiantian Lu, Jiuyu Jin, Tingting Wang, Yajing Zhang, Yue Dou, Morihisa Fujita, Yixuan Xie, Yishi Liu, Ran Xie

## Abstract

GlycoRNAs, small non-coding RNAs covalently linked to *N*-glycans, reveal an unexpected intersection between RNA biology and glycosylation, with potential roles in immunity and cell-cell communication. The origins and biogenesis of their glycans remain unclear. Using CRISPR-Cas9 screening, metabolic glycan labeling and lysosome-targeted proteomics, we reconfirm canonical *N*-glycosylation enzymes (STT3A/B) and identify components of the mannose-6-phosphate (M6P) lysosomal sorting pathway (GNPTAB, M6PR) as essential contributors. Strikingly, lysosomal aspartylglucosaminidase (AGA) acts as a rate-limiting regulator, linking glycoprotein catabolism to glycoRNA formation. Loss of AGA collapses both abundance and diversity of RNA-linked *N*-glycans, while pulse-chase tracing shows glycoprotein-derived glycans are repurposed for RNA glycosylation. These findings position lysosomes as active participants in RNA modification and redefine glycoRNA biogenesis as a metabolically integrated process.

## Introduction

Glycosylated RNAs (glycoRNAs) have emerged as an unexpected class of RNA-containing glycoconjugates that expand the molecular vocabulary of the cell surface.^1^ Composed of small noncoding RNAs bearing complex glycans, most prominently sialylated *N*-glycans,^1^ glycoRNAs extend glycosylation beyond proteins and lipids and implicate RNA within glycan-dependent processes such as immune recognition, cargo trafficking, and cell-cell communication.^2–5^ However, this field remains mechanistically nascent. Beyond the identification of acp^3^U as a glycan-linked RNA nucleoside, the origin of glycoRNA glycans,^6^ the chemical basis of glycosidic bond, the pathways that assemble and remodel glycoRNAs, and their coordination with cellular homeostasis remain largely unresolved.^7^

Canonical *de novo N-glycosylation* offers a framework for understanding glycoRNA biogenesis.^8^ In this pathway, lipid-linked oligosaccharides are assembled in the ER, transferred to nascent substrates by oligosaccharyltransferase (OST) complex, remodeled it the Golgi into high-mannose, hybrid, and complex *N*-glycans, and ultimately trafficked to the cell surface, secreted, or *en route* endolysosomal compartments for turnover and recycling (**Figure 1, steps 1-4**). Consistent with this framework, perturbation of GALE (UDP-galactose rheostat enzyme) or STT3A/3B (catalytic subunits of mammalian OST) activity, *ldlD*-dependent glycan metabolism, or downstream *N*-glycan processing impairs glycoRNA formation,^1,9,10^ supporting a requirement for the canonical *N*-glycosylation pathway (**Figure 1**, **middle**). However, *N*-glycan homeostasis depends on more than biosynthesis: lysosomal turnover liberates reusable sugars, ***salvage pathways*** rebuild nucleotide-sugar donors, and the ***mannose-6-phosphate (M6P) lysosomal sorting pathway*** delivers the hydrolases that sustain this catabolic arm (**Figure 1**, **step 4)**.^11^ This endolysosomal axis also intersects with RNA biology, as SIDT1/2-mediated RNA transport links RNA trafficking to lysosomal compartments,^2^ whereas DTWD2 may license selected RNAs for ER-associated glycoRNA production (**Figure 1 bottom left**).^6^ These connections position glycoRNAs within the broader glycoconjugate economy of the cell and raise the central question: how are glycoprotein *N*-glycan turnover and glycoRNA homeostasis mechanistically bridged?

**Figure 1.**
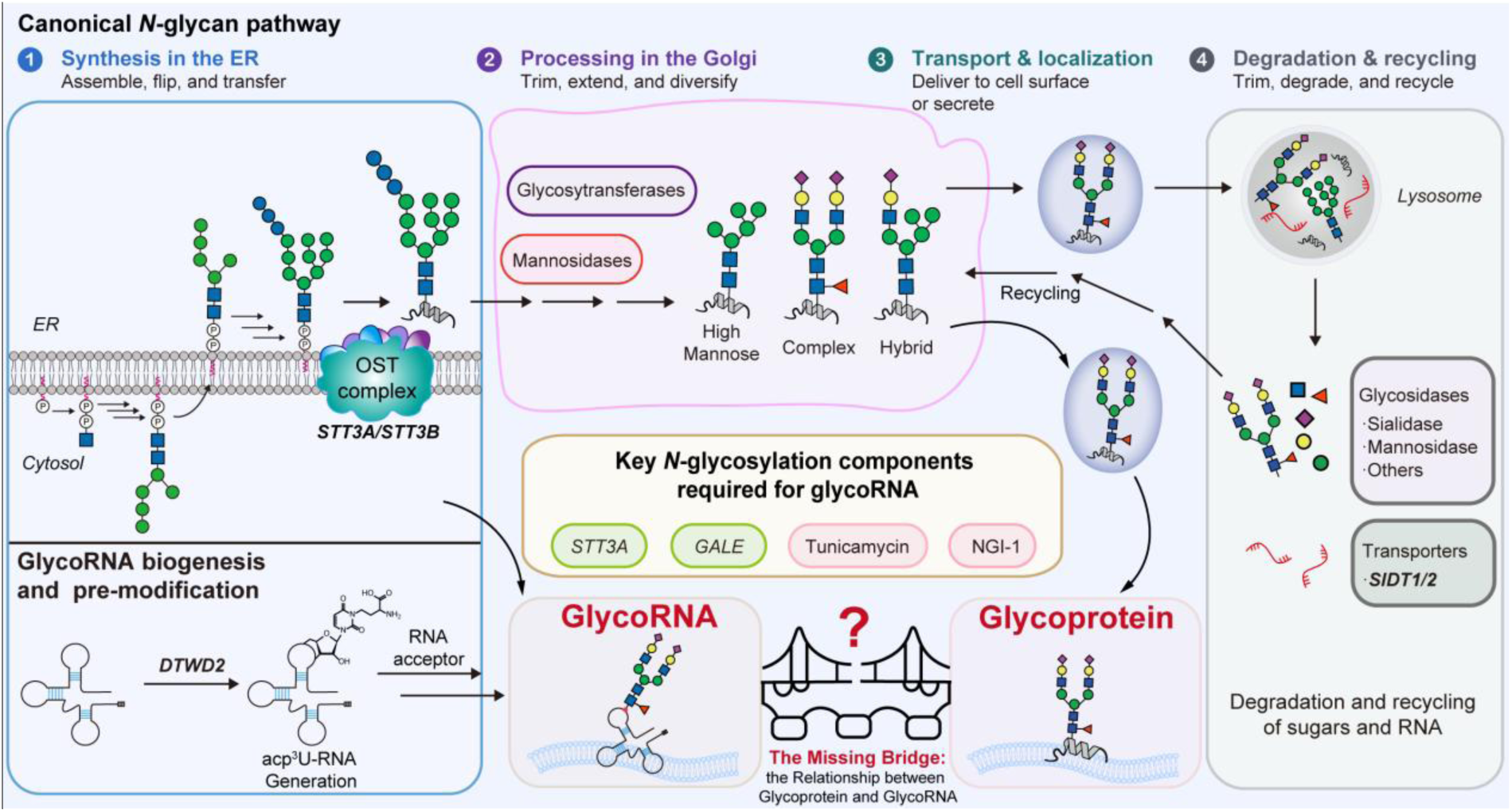
Proposed connection between canonical N-glycan metabolism and glycoRNA biogenesis. Schematic summary of *N*-glycan synthesis, processing, trafficking, and turnover in relation to glycoRNA formation. In the canonical pathway, lipid-linked oligosaccharides are assembled in the ER and transferred by the OST complex, including **STT3A/STT3B**, followed by Golgi processing to generate *N*-glycans. Mature glycoconjugates are transported to the cell surface, secreted, recycled, or degraded in lysosomes by glycosidases, with **SIDT1/2** implicated in RNA/glycoconjugate transport and turnover. The lower panel highlights proposed glycoRNA biogenesis, including **DTWD2**-dependent RNA pre-modification and *N*-glycosylation-related regulators (**STT3A**, **GALE**, tunicamycin, and NGI-1). The central unresolved question is the “missing bridge” connecting glycoprotein and glycoRNA pathways, including whether they share enzymes, trafficking routes, and degradation mechanisms.

Herein, using CRISPR-Cas9 screening, metabolic glycan labeling (MGL), and organelle-resolved proteomics, we identify the lysosomal aspartylglucosaminidase (AGA) as a pivotal upstream regulator of glycoRNA production. We show that lysosomal glycoprotein catabolism fuels glycoRNA synthesis, and delineate how *de novo* and salvage pathways converge to support this process. the interplay between *de novo* and salvage pathways. Together, these findings establish a mechanistic framework for glycoRNA biogenesis and unravel lysosomes as unexpected contributors to RNA glycosylation.

## Results

### Platform for GlycoRNA Genetic Screening

To elucidate the genetic basis of glycoRNA biogenesis, we leveraged a previously generated stable knockout (KO) library targeting 36 core *N*-glycosylation genes based on HEK293 cells,^10,12^ spanning *de novo* biosynthesis, lysosomal sorting, and salvage pathways (**Table S1**). Sialylated glycoRNAs were quantitatively assayed with a recombinant Siglec11-IgG Fc fusion protein (Siglec11-Fc; **Figure S1A**),^1,5,10^ whose binding was strongly reduced by RNase and colocalized with *Maackia amurensis* lectin II (MAL-II; α2,3-sialic acid) and *Sambucus nigra* lectin (SNL; α2,6-sialic acid), confirming cell-surface sialoglycoRNA binding affinity of Siglec11-Fc (**Figures S1B-S1E**). Assay performance was validated using SIDT2- and STT3A-deficient cells, which showed the expected reduction in Siglec11-Fc signal by flow cytometry (**Figures S1F-1H**).^1,2,10^ In parallel, three *N*-azidoacetylmannosamine derivatives (**Figure 1I**) were used to hijack the sialic acid salvage pathway,^13–15^ enabling azide incorporation into sialylated glycoRNAs and subsequent detection via bioorthogonal ligation with dibenzocyclooctyne-biotin (DBCO-biotin).^16^ All analogs labeled glycoRNAs reproducibly (**Figure S1J**), and the most accessible peracetylated *N*-azidoacetylmannosamine (Ac_4_ManNAz) was selected to validate screening results.

### Lysosomal Pathways Drive Sialylated GlycoRNA Biogenesis

To orthogonally validate the genetic screening via the KO library, we quantified sialylated glycoRNAs by anti-biotin Northern blot in cells lacking the core *N*-glycosylation enzymes STT3A/B. Consistent with prior work, STT3A/B disruption markedly reduced glycoRNA levels (**Figure S1K**), confirming dependence on the *de novo N*-glycosylation pathway.^10^ Unexpectedly, components of the M6P lysosomal sorting pathway were also required:^11^ loss of *GNPTAB* or *M6PR* diminished Siglec11-Fc binding and glycoRNA signals (**Figures 2A-2C**), linking lysosomal trafficking to glycoRNA biogenesis. In contrast to impaired sialylated *N*-glycan formation, parallel wheat germ agglutinin (WGA) staining showed minimal changes, highlighting the specific RNase sensitivity of Siglec-11 binding**(Figures 2D-2E**). Although M6PR loss produced weaker effects in flow cytometry than in Northern blotting (**Figure 2F**), likely due to redundancy with the cation-independent M6PR (IGF2R),^17^ we reasoned that genuine glycoRNA regulators should confer increased RNase sensitivity due to reduced cell-surface glycoRNAs. The weaker *GNPTAB* phenotype likely reflects partial loss in bulk populations, as complete deficiency is lethal (**Figure 2F**).^18^ Together, these results identify M6P-mediated lysosomal sorting as an unexpected but essential determinant of sialylated glycoRNA biogenesis, acting by sustaining cellular sialylation capacity through glycan turnover and salvage.

**Figure 2.**
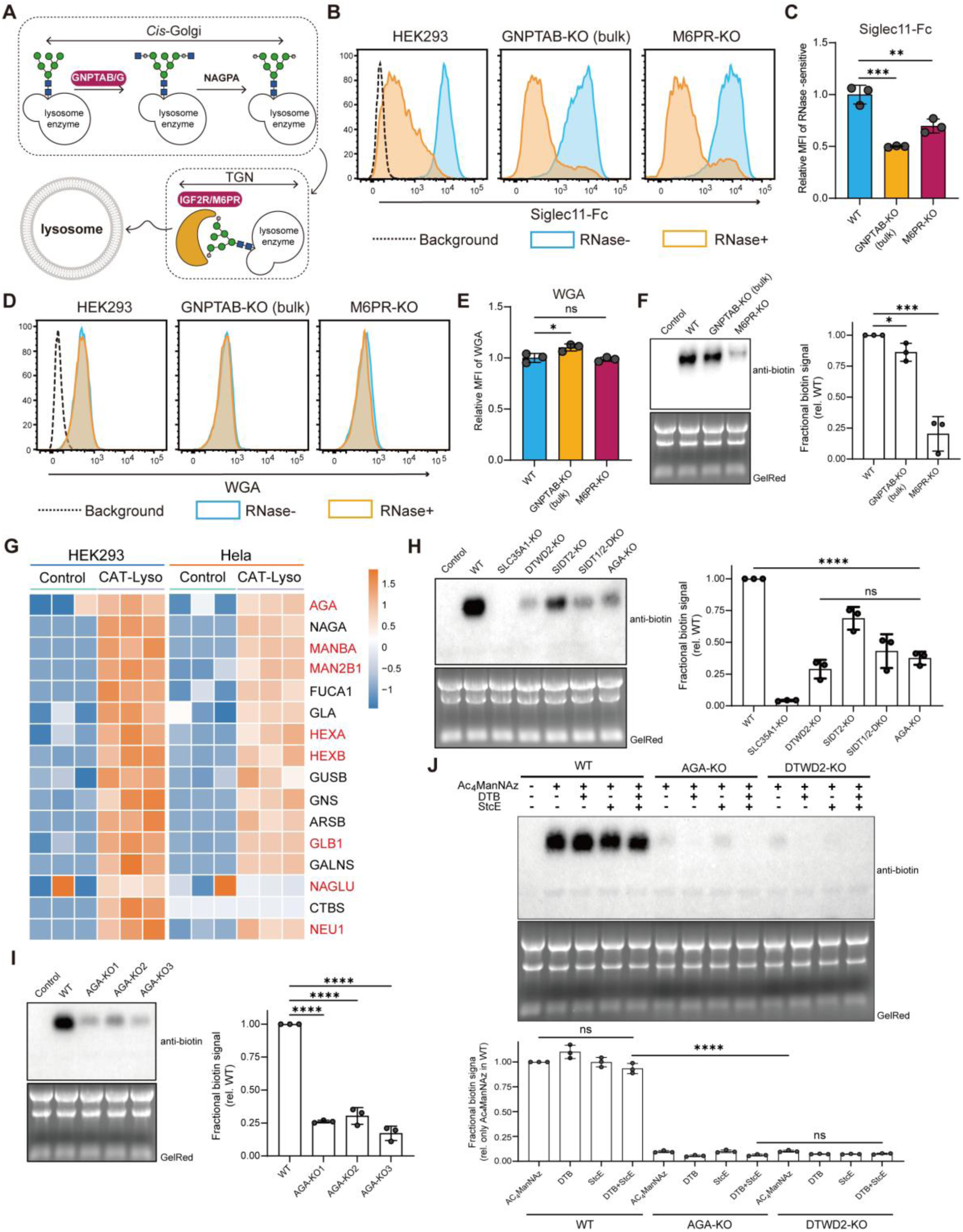
Impact of lysosomal glycosidases on glycoRNA levels. **(A)** Diagram of the mannose-6-phosphate (M6P) lysosomal sorting pathway. **(B)** Flow cytometry histograms of GNPTAB-KO or M6PR-KO cells stained with Siglec11-Fc. **(C)** Quantification of flow cytometry data in (A). The relative level of MFI in WT was set to 1, and GNPTAB-KO and M6PR-KO levels were displayed as the mean ± SD of three independent experiments. **(D)** Flow-cytometry analysis of WGA staining in HEK293, GNPTAB-KO bulk, and M6PR-KO cells with or without RNase treatment. **(E)** Quantification of flow cytometry data in (D). The relative level of MFI in WT was set to 1, and GNPTAB-KO and M6PR-KO levels were displayed as the mean ± SD of three independent experiments. **(F)** Northern blotting of RNA from HEK 293 wild-type (WT). GNPTAB-KO or M6PR-KO cells treated with 100 mM Ac_4_ManNAz for 48h. After RNA purification, azide on glycoRNA was conjugated to DBCO-biotin, and visualized with streptavidin-HRP. Total RNA was stained and imaged with GelRed for data quality interrogation. Inset: quantification of the blot in (F) from biological triplicates. **(G)** Heatmap plot of Z scores of the lysosomal hydrolases abundances from CAT-Lyso. **(H)** Northern blotting of RNA from HEK 293 WT. SLC35A1-, DTWD2-, SIDT2-. SIDT1/2- or AGA-KO cells treated with 100 mM Ac_4_ManNAz for 48h. Inset: quantification of the blot in (H) from biological triplicates. **(I)** Northern blotting of RNA extracted from HEK 293 WT or triplicate biological AGA-knockout clones treated with 100 mM Ac_4_ManNAz for 48h. Inset: quantification of the blot in (I) from biological triplicates. **(J)** Northern blotting of RNA extracted from HEK 293 WT, DTWD2-KO or AGA-KO clone treated with 100 mM Ac_4_ManNAz for 48h, which were subjected to the conventional nondenaturing proteinase K treatment or to the modified proteinase K treatment in DTB, and then further treated with mucinase StcE protease. Inset: quantification of the blot in (J) from biological triplicates. Data represent mean ± SEM of three independent experiments; statistical significance was determined by two-tailed Student’s *t*-test: ns, not significant, \**P* < 0.05, \*\**P* ≤ 0.01, \*\*\**P* ≤ 0.001, \*\*\*\**P* ≤ 0.0001.

### Lysosomal Proteomics Implicates AGA in GlycoRNA Biogenesis

Guided by M6P pathway findings and SIDT knockout effects,^2^ we mapped lysosomal contributions to glycoRNA biogenesis using CAT-Lyso,^19^ a lysosome-targeted photocatalytic labeling approach, to map the proteome *in situ* in HEK293 and HeLa cells (**Figure 2G, S2A and S2B**). Mass spectrometry (MS) and Gene Ontology (GO) analysis identified 1041 and 1099 proteins, respectively (**Figures S2C, Table S2**), with strong enrichment of lysosomal components, including 14 glycosidases involved in *N*-glycan salvage, such as endo-β-*N*-acetylglucosaminidase (NAGLU), α-mannosidases (MAN2B1/MANBA), β-hexosaminidases (HEXA/HEXB), β-galactosidase (GLB1), and α-neuraminidase (NEU1) (**Figure 2F, S2D and S2E**), highlighting lysosomal enzymes as key glycoRNA modulators.

We then focus on lysosomal aspartylglucosaminidase *N*(4)-(β-*N*-acetylglucosaminyl)-L-asparaginase (EC 3.5.1.26, AGA), which release free *N*-glycans from glycoproteins for glycan salvage.^20^ Intriguingly, AGA loss sharply reduced cell-surface sialylated glycoRNAs, as assessed by Ac_4_ManNAz labeling, with a substantially stronger effect compared to knockouts of known glycoRNA regulators SLC35A1,^21^ DTWD2, or SIDT1/2 (**Figure 2H**). This phenotype was robust and reproducible across three independent AGA knockout clones (**Figure 2I**), revealing an unexpected yet essential role for AGA in glycoRNA biogenesis and stability. Independent validation using the RNA-optimized periodate oxidation and aldehyde labeling (rPAL) assay^6^ likewise confirmed the loss of sialylated glycoRNAs in AGA-deficient cells (**Figure S2F**), including under an RNase-normalized analysis framework (**Figure S1G and S2G**). To address potential RNase-insensitive contamination during glycoRNA isolation,^22,23^ we introduced protease-based controls in WT, AGA-KO and DTWD2-KO HEK293 cells, followed by MGL and proteomic analysis **(Figure 2J, S2H-S2I and Table S3)**. These results indicate that (glyco)protein contamination does not explain the observed signal, supporting the ability of the assay to resolve bona fide glycoRNA species from co-purified contaminants.

To capture glycoform-level changes comprehensively, we profiled RNA-linked *N*-glycans using GlycanDIA,^24,25^ a data-independent acquisition-based quantitative glycomics platform. Small RNAs were purified, enzymatically stripped of protein contaminants, and *N*-glycans were released by PNGase F, enriched by porous graphitized carbon solid-phase extraction (PGC-SPE), and analyzed by mass spectrometry (**Figure 3A**). AGA knockout caused a global collapse in both abundance and diversity of glycoRNA-associated *N*-glycans, with coordinated losses across different glycan subtypes, including high mannose, undecorated, fucosylated, sialylated, and sialofucosylated glycans (containing both fucose and sialic acid), mirroring our sialoglycoRNA blotting results using Ac_4_ManNAz and rPAL. **(Figures 3B, 2H, and S2F)**. DTWD2-KO cells, which lack the RNA *N*-glycosylation site, displayed an indistinguishable glycomic profile, also consistent with MGL assays (**Figures 3C-3D and 2J**). The near-identical phenotypes establish AGA as a rate-limiting enzyme for glycoRNA biogenesis, directly linking lysosomal *N*-glycan salvage pathway to glycosylated RNA production. SWAMNA-based nucleoside analysis^26^ further showed that acp^3^U, the DTWD2-installed uridine modification proposed to serve as the glycan acceptor, is depleted in DTWD2-KO cells but accumulates in AGA-KO cells in both total and membrane-associated RNA (**Figures 3E-3G**). Thus, AGA loss appears to impair glycosylation downstream of acceptor-site installation rather than eliminating the RNA substrate itself.

**Figure 3.**
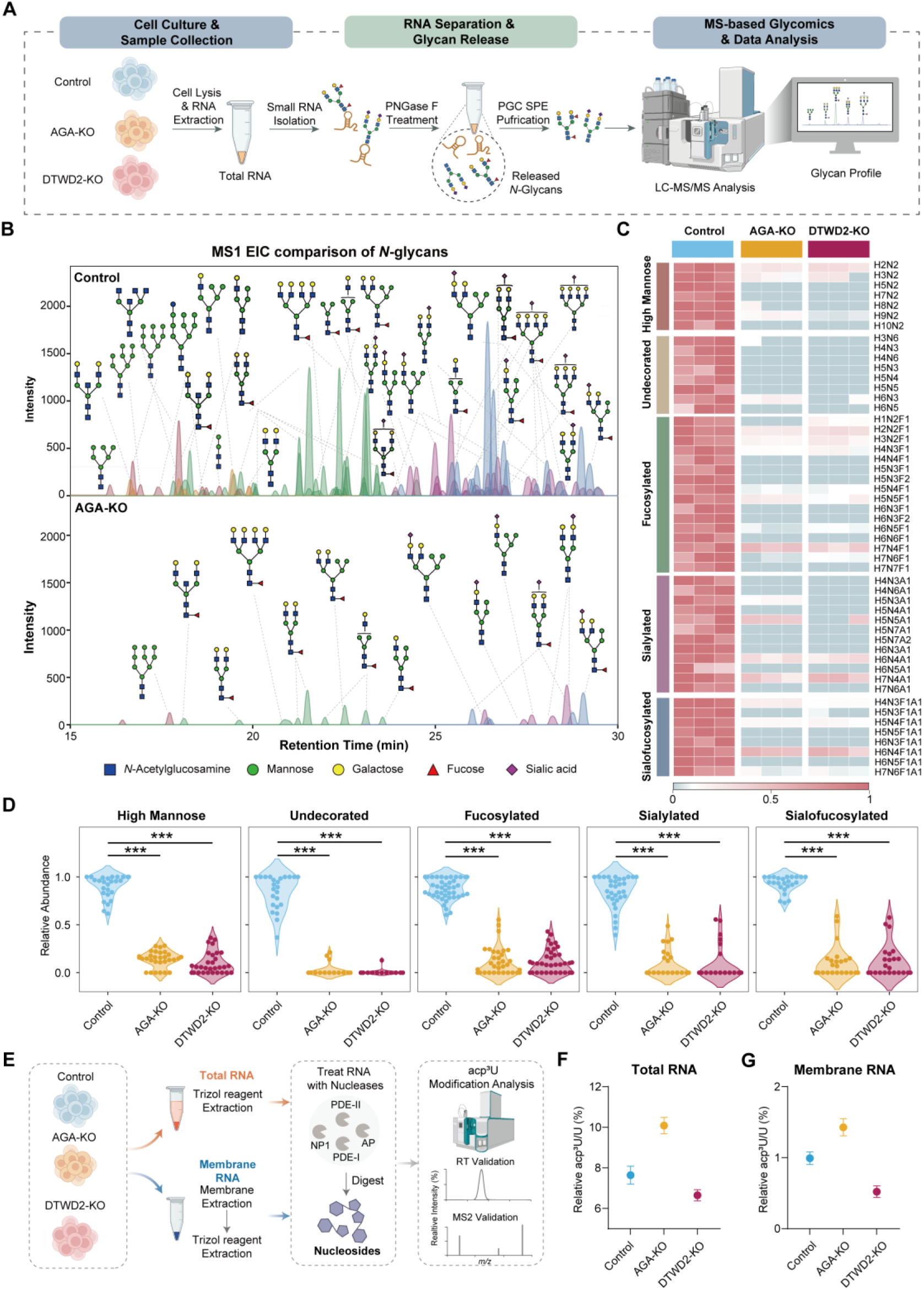
GlycanDIA reveals RNA *N*-glycan remodeling in HEK293 cells. **(A)** Schematic of the GlycanDIA workflow for releasing glycans from RNA and purifying free glycans for mass spectrometry analysis. **(B)** Representative chromatograms comparing *N*-glycome profiles of HEK293 WT and AGA-KO cells. **(C)** Heatmap of total RNA *N*-glycans identified in HEK293 WT, DTWD2-KO, and AGA-KO cells, showing distinct glycan profiles across samples. **(D)** Relative abundance of five glycan subtypes in glycoRNAs from HEK293 WT, DTWD2-KO, and AGA-KO cells. **(E)** Schematic of the acp^3^U modification analysis from total RNA and membrane RNA. **(F, G)** Quantification of normalized acp^3^U levels from small RNAs of total RNA (F) and membrane RNA (G) isolated from HEK293 WT, DTWD2-KO, and AGA-KO cells by SWATH-MS. Data represent mean ± SEM of three independent experiments; statistical significance was determined by two-tailed Student’s *t*-test: \*\*\**P* ≤ 0.001.

### Biochemical Validation of AGA *N*-Glycan-Releasing Activity

AGA is a lysosomal *N*-terminal nucleophile hydrolase that cleaves the GlcNAc-Asn bond of *N*-glycoproteins.^27–29^ *In vitro*, purified AGA efficiently hydrolyzed GlcNAc-Asn and a complex sialylated glycopeptide (SGP), releasing *N*-glycans from native cellular glycoproteins at levels comparable to PNGase F (**Figures 4A-4C**).^30^ In striking contrast, AGA exhibited minimal activity toward RNA-linked *N*-glycans (**Figure 4C**), demonstrating a strong intrinsic preference for glycoproteins over glycoRNAs. These data demonstrate that AGA functions as a legitimate lysosomal *N*-glycan releasing enzyme but does not directly deglycosylate glycoRNAs, implying that its impact on glycoRNA abundance is indirect and mediated through glycan salvage rather than direct RNA deglycosylation.

**Figure 4.**
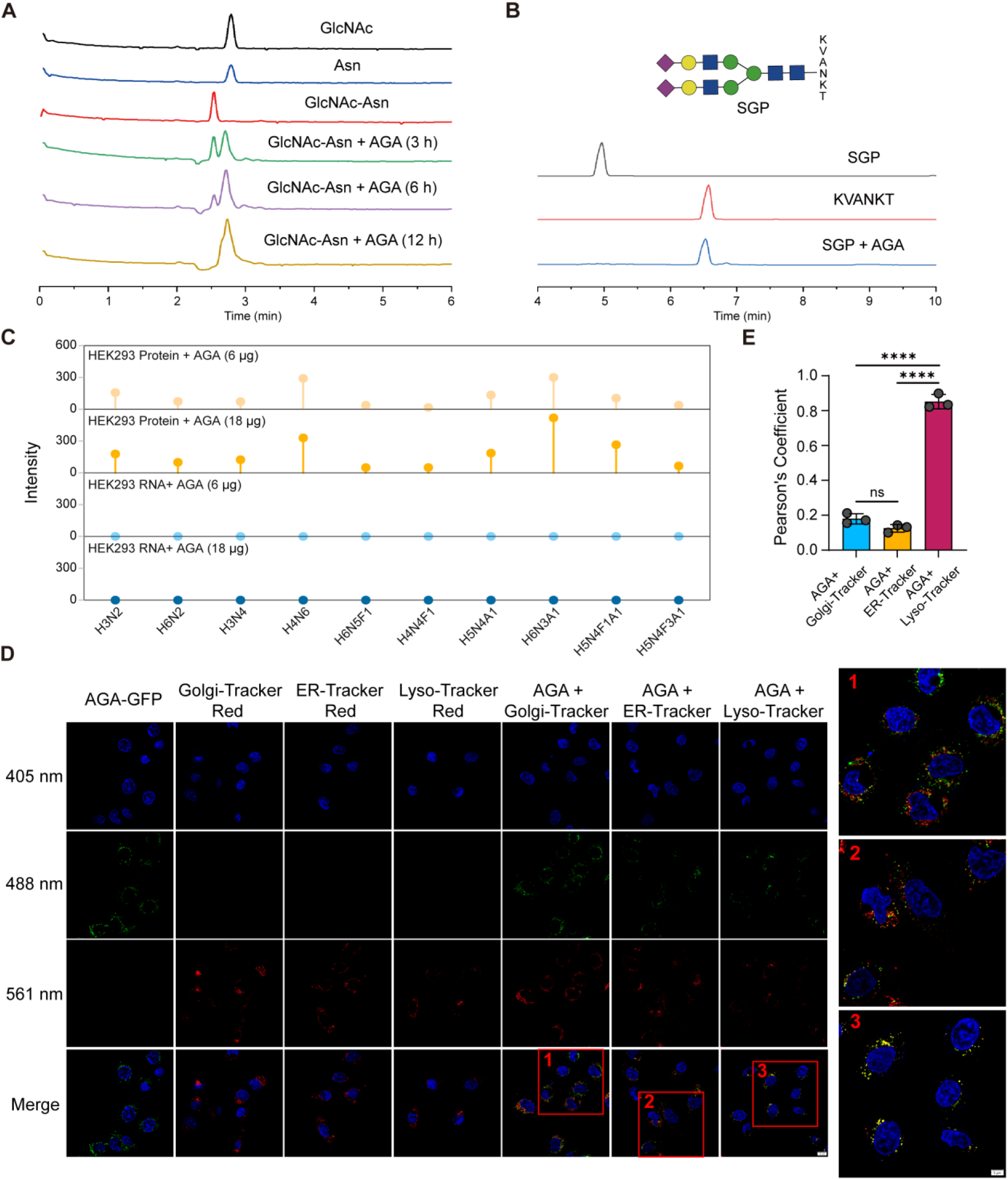
Subcellular localization and enzymatic activity of AGA in living cells. **(A)** LC-MS analysis of enzymatic products from GlcNAc-Asn incubated with 25 μL AGA in assay buffer (50 mM sodium phosphate pH 6.0, 150 mM NaCl, 5 mM DTT) at 37 °C for indicated times. Products were desalted and analyzed by LC-MS. **(B)** HPLC analysis of products from SGP incubated with 25 μL AGA in assay buffer at 37 °C for 12 h. Then products were desalted and analyzed byHPLC. **(C)** GlycanDIA analysis of representative glycans released from HEK293 proteins or sRNA treated with 6 μg or 18 μg AGA, respectively. **(D)** Representative confocal images of AGA-GFP (FITC channel, λex = 488 nm) in living cells. Cells were co-stained with Golgi-, ER-, or Lyso-Tracker Red to indicate organelles, and nuclei were stained with Hoechst 33342. Scale bars: 10 μm (main), 5 μm (zoomed merge). **(E)** Pearson’s correlation coefficients quantifying colocalization of AGA-GFP with organelle markers. Data represent mean ± SEM of three independent experiments; statistical significance was assessed by two-tailed Student’s t-test: ns, not significant, \*\*\*\**P* ≤ 0.0001.

### AGA Structure-Function Coupling Links *N*-Glycan Salvage to GlycoRNA Biogenesis

We next asked how AGA’s lysosomal function and disease-relevant catalytic architecture couple *N*-glycan salvage to glycoRNA biogenesis in cells. Loss of AGA causes the lysosomal storage disorder aspartylglucosaminuria (AGU),^27,28^ highlighting its essential role in *N*-glycoprotein turnover. Consistent with its role in lysosomal catabolism, GFP-tagged AGA localized predominantly to lysosomes, co-localizing with LysoTracker but not ER or Golgi markers, with weak non-lysosomal signals likely reflecting transient trafficking through the M6P sorting pathway (**Figure 4D and 4E**). Functionally, AGA knockout caused a pronounced loss of sialylated glycoRNAs, whereas AGA complementation partially restored glycoRNA sialylation, as assessed by azidosugar labeling, rPAL analysis, and GlycanDIA profiling (**Figures S3**).

AGA is produced as a 346-aa zymogen that undergoes autocatalytic cleavage at D205-T206 to generate an active (αβ)_2_ heterotetramer. Structural analyses define a funnel-shaped active site organized around the *N*-terminal nucleophile T206, with C306 required for proper folding, R234 and D237 mediating substrate engagement, and AGU-associated mutations such as T257I disrupting catalytic efficiency or substrate affinity.^29,31–33^ Guided by these structural and clinical insights, we selected mutations targeting catalytic, structural, and disease-linked residues to dissect how impaired *N*-glycan release impacts glycoRNA production (**Figure S4A**). Mutations in key catalytic or structural residues uniformly suppressed glycoRNA abundance (**Figures S4B and S4C**), despite mutation-specific alterations in glycoRNA glycoform composition (**Figure S4D and S4E**), indicating that intact AGA activity is required to sustain glycoRNA production. In contrast, AGA deficiency led to AGU-like accumulation of *N*-glycans on total glycoproteins **(Figure S5A and S5B)**, with elevated cell-surface glycosylation confirmed by lectin profiling **(Figure S5C).** Consistently, disruption of *de novo N*-glycosylation by treating cells with brefeldin A (BFA),^34^ swainsonine (SW)^35^ or tunicamycin (TU)^36^ reduced glycoRNA abundance **(Figure S6A-S6C)**, supporting a model in which lysosomal salvage and *de novo* synthesis jointly sustain glycoRNA biogenesis.

### GlycoRNA Biogenesis Is Coupled to Glycoprotein *N*-Glycan Turnover

To directly test whether lysosomal *N*-glycan catabolism from glycoproteins fuels glycoRNA biogenesis, we performed classical pulse-chase metabolic tracing using ^13^C-glucose. HEK293 cells were pulsed with ^13^C-glucose and chased for up to 48 h, followed by MS analysis of *N*-glycans from glycoproteins and glycoRNAs (**Figure 5A**). Quantitative isotope incorporation revealed a clear temporal hierarchy (**Figure 5B**): For the representative *N*-glycan subtypes, namely high mannose H9N2, complex H5N4F1A1 and hybrid H6N3, ^13^C-labeled *N*-glycans appeared first in glycoproteins and only later in glycoRNAs, indicating a lag phase in glycoRNA labeling (**Figure 5C**). Isotope incorporation into glycoRNA reached only ∼50% of that observed in glycoproteins at matched time points within 24 h **(Figure 5D)**, inconsistent with a model in which *de novo* nucleotide-sugar biosynthesis supplies all glycoconjugates simultaneously. To further examine glycan reutilization, we performed a reverse pulse-chase experiment: cells were first labeled with ^13^C-glucose for 24 h, then chased with unlabeled ^12^C-glucose **(Figure 5E).** Our results revealed faster ^13^C signal decay in glycoprotein-derived *N*-glycans than in glycoRNAs, again with a pronounced lag **(Figure 5F and 5G)**. Analysis of isotopologues (*e.g.,* [M+53], [M+67]) in representative H8N2 and H5N4F1A1 sugar chains confirmed delayed turnover and prolonged retention in glycoRNAs relative to glycoproteins. Notably, labeled glycoRNA signals transiently increased during early chase phases before declining, consistent with continued glycoRNA assembly from pre-existing glycan pools. Together, these pulse-chase experiments provide direct kinetic evidence that glycoRNA *N*-glycans are supplied, at least in part, by recycled glycoprotein-derived glycans rather than exclusively by *de novo* biosynthesis, establishing lysosomal glycan salvage as a functional upstream source for glycoRNA biogenesis.

**Figure 5.**
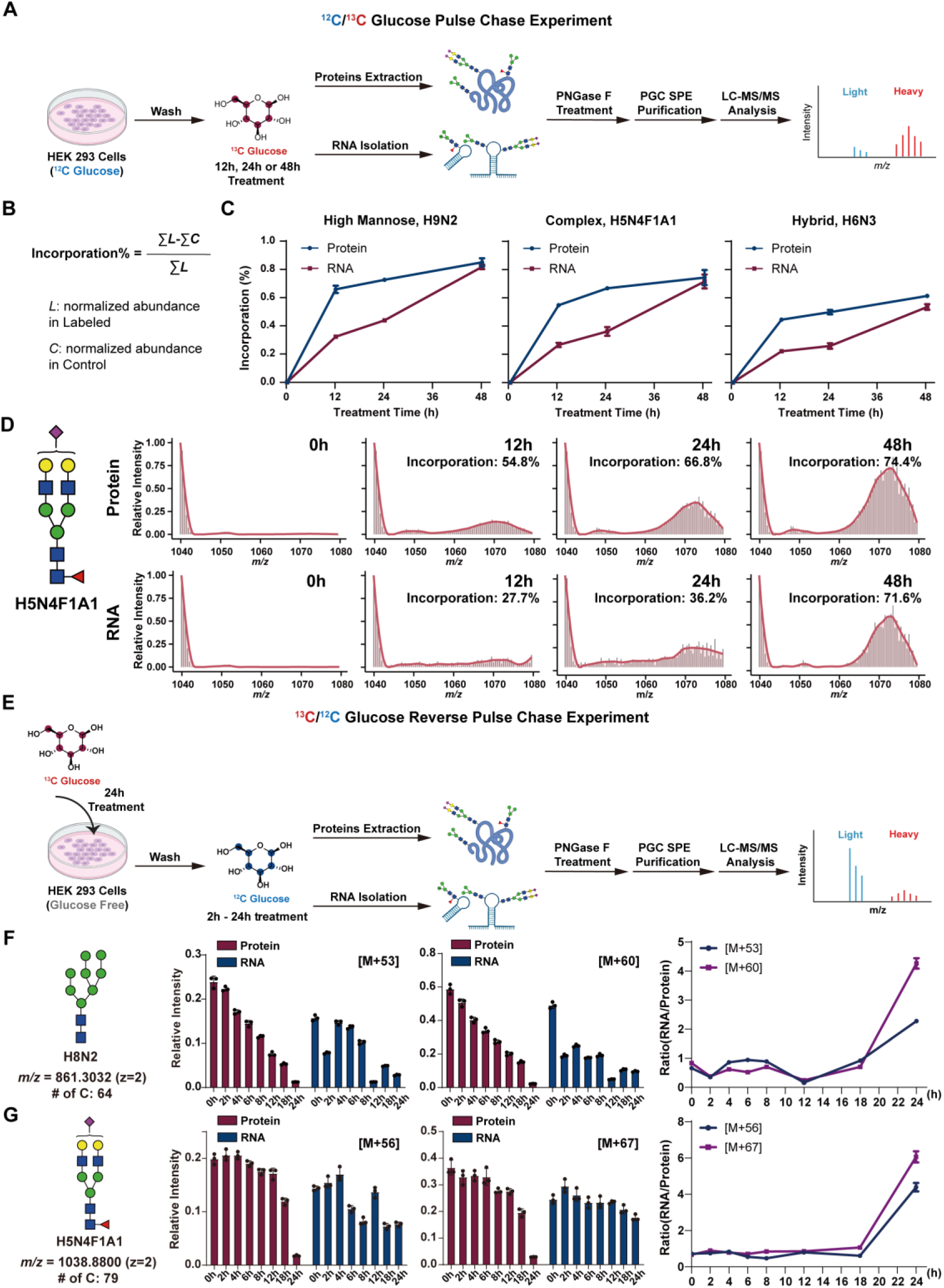
Metabolic pulse-chase of *N*-Glycans in Protein and RNA. **(A)** Workflow of the ^12^C/^13^C-glucose pulse-chase experiment. HEK293 cells at 50% confluence were cultured in glucose-free DMEM supplemented with ^13^C glucose (4.5 g L^-1^) for 12, 24, or 48 h. Cells were harvested for protein extraction and RNA isolation, and ^13^C incorporation into glycan monosaccharides was quantified by LC-MS/MS. **(B)** Quantitation using summed normalized abundances of each compound from labeled and unlabeled samples. **(C)** Rates of ^13^C incorporation into various *N*-glycan types over time. **(D)** Representative MS spectra and ^13^C-incorporation analysis of *N*-glycan H5N4F1A1 from protein or RNA after 12–24 h of ^13^C glucose treatment. **(E)** Workflow of ^13^C/^12^C glucose reverse pulse-chase experiment. After 24 h of ^13^C glucose, cells were washed and cultured in ^12^C glucose for indicated times before harvesting for protein and RNA analysis of glycan ^13^C content. **(F, G)** MS spectra of *N*-glycan H8N2 (F) and H5N4F1A1 (G) from protein or RNA after 2-24 h of ^12^C glucose chase.

### Dynamic AGA Interactome Links Trafficking to GlycoRNA Biogenesis

Finally, to map the AGA interactome, we transfected FLAG-tagged AGA (**Figure 6A**) and performed immunoprecipitation followed by mass spectrometry. A total of 169 candidate interactors were identified, with proteins involved in lysosomal trafficking (AP3B1, AP3S1, RAB38),^37,38^ Golgi/retrograde transport (COG2-5, COG7/8, COPB2, VPS53),^39,40^ and ER-Golgi cargo transport (LMAN1, YIF1A)^41,42^ showing significantly elevated spectra counts relative to controls (**Figure 6B and Table S4**). This broader AGA-centered network suggests interactions extend beyond lysosomal proteins (**Figure 6C**), highlighting dynamic trafficking pathways that may pinpoint the actual sites of glycoRNA synthesis, not necessarily restricted to lysosomes. Overlaps between CAT-lyso and AGA-IP datasets further support this dynamic, remote interaction model (**Figure 6D**). Consistently, RNA sequencing of isolated lysosomes revealed glycosylated donor RNAs, including tRNAs and Y RNAs, providing mechanistic evidence for potential substrate supply (**Figure 6E**). Collectively, these findings indicate that the glycoRNA landscape is shaped by unexpected, dynamic AGA interactions, underscoring substantial unexplored biology in glycoRNA research.

**Figure 6.**
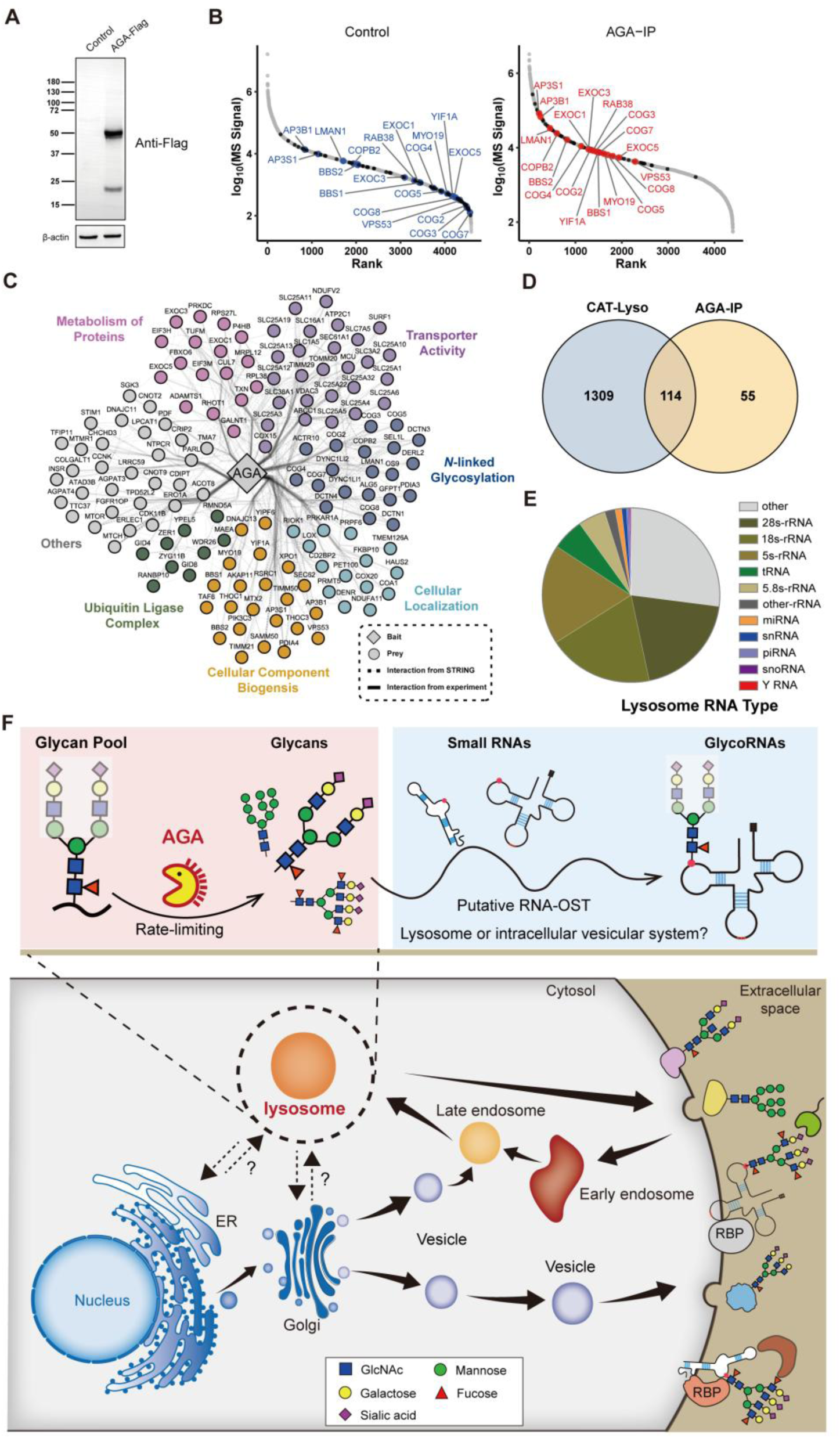
Identification of AGA-interacting proteins and lysosomal RNAs. **(A)** Western blot of HEK293 cell lysates transfected with Flag-tagged AGA. **(B)** Mass spectrometry analysis of proteins interacting with AGA. **(C)** Protein interaction network from AGA-Flag immunoprecipitation, covering 169 proteins. **(D)** Venn diagram comparing CAT-Lyso and AGA-IP protein datasets. **(E)** Analysis of lysosomal RNAs associated with AGA. **(F)** Proposed model for lysosomal glycoprotein catabolism-coupled glycoRNA biogenesis. Canonical ER-Golgi *N*-glycosylation generates glycoproteins that traffic through the cell surface and endolysosomal system. In lysosomes, **AGA** releases reusable *N*-glycan pools from catabolized glycoproteins, which are proposed to support small RNA glycosylation through an unidentified OST-like activity and vesicular trafficking intermediates. AGA loss depletes glycoRNA abundance and glycan diversity while increasing glycoprotein *N*-glycans, supporting lysosomal glycan salvage as an upstream source for glycoRNA production.

## Discussion

Our study identifies glycoRNA biogenesis as a system-level process that integrates canonical *N*-glycosylation, lysosomal sorting, and glycan salvage pathways (**Figure 1**). While previous work hints the requirement for *de novo N*-glycosylation enzymes in glycoRNA formation, our genetic screen unexpectedly uncovered a central role for lysosomal function, particularly the M6P-dependent sorting machinery. Disruption of GNPTAB or M6PR impaired sialylated glycoRNA expression to a degree comparable to loss of core biosynthetic enzymes, demonstrating that intact lysosomal trafficking is essential for sustaining glycoRNA production. These findings expand the conceptual framework of glycoRNA biogenesis beyond Golgi-centric biosynthesis and place lysosomal integrity as a critical upstream determinant of surface glycoRNA abundance.

A key mechanistic insight of this study is that AGA acts as a rate-limiting regulator of glycoRNA biogenesis (**Figure 6F**). Traditionally, AGA is considered a terminal lysosomal hydrolase essential for glycoprotein degradation and linked to disease. Using orthogonal detection approaches, our data reveal an unanticipated role for AGA in providing glycan substrates for glycoRNA formation, with observed signals primarily reflecting glycoRNAs rather than glycoproteins. AGA deficiency phenocopied loss of the RNA glycosylation site itself (DTWD2-KO), collapsing both the abundance and structural diversity of RNA-linked *N*-glycans, despite causing accumulation of *N*-glycans on glycoproteins. Similarly, knockout of the lysosomal RNA transporter SIDT caused a comparable decrease in glycoRNA, consistent with previous reports.^2^

Importantly, AGA exhibited strong selectivity for glycoprotein-linked *N*-glycans and negligible activity toward glycoRNAs, indicating that its role in glycoRNA biogenesis is indirect, *i.e.,* acting upstream to liberate reusable glycan pools rather than remodeling glycoRNAs directly. The dual phenotype, *i.e.,* glycoprotein hyper-glycosylation alongside glycoRNA loss, positions AGA as a metabolic gatekeeper balancing glycan degradation with noncanonical RNA glycosylation. Complementation suggests additional factors, as most RNA glycans are not restored and glycoform-specific effects persist.

Metabolic pulse-chase tracing provides direct kinetic evidence that glycoRNA *N*-glycans are supplied, at least in part, by recycled glycoprotein-derived glycans. Isotope incorporation followed a clear temporal hierarchy, with labeled *N*-glycans appearing first on glycoproteins and only later on glycoRNAs. Reverse pulse-chase experiments further demonstrated faster isotopic decay in glycoproteins than in glycoRNAs, revealing delayed turnover and prolonged retention of glycans on RNA. Together, these data support a model in which lysosomal degradation of glycoproteins generates a salvageable glycan pool that is subsequently repurposed for glycoRNA assembly. This coupling of catabolism and biosynthesis establishes glycoRNA biogenesis as a metabolically integrated process and suggests that RNA glycosylation may act as a downstream sensor of cellular glycan flux, lysosomal health, and glycoprotein turnover.

Although AGA is a soluble lysosomal luminal enzyme, its interactome was unexpectedly enriched for cytosolic trafficking factors, including AP3 and COG complex components. These associations are unlikely to reflect direct physical interactions, but instead likely arise from indirect or complex-mediated coupling through membrane-associated intermediates, such as transmembrane receptors or sorting machinery engaged during endolysosomal trafficking. Consistent with this interpretation, overlap between CAT-Lyso and AGA-IP datasets suggests that

AGA associates with dynamic trafficking hubs rather than acting exclusively within mature lysosomes. This raises the possibility that glycoRNA synthesis or modification occurs along the endolysosomal-Golgi continuum, rather than being confined to terminal lysosomes. Supporting this model, glycosylated donor RNAs, including tRNAs and Y RNAs, were detected in isolated lysosomes, indicating that relevant RNA substrates are present within these compartments.^43^

### Limitations of the study

Regrettably, no single assay or binder at present can unambiguously and comprehensively quantify glycoRNAs; instead, their detection relies on convergent evidence from metabolic glycan labeling, enzymatic sensitivity, genetic perturbation, and orthogonal biochemical readouts. Consequently, the field still lacks state-of-art tools to selectively detect glycoRNAs with defined *N*-glycan structures, likely contributing to discrepancies among rPAL, GlycanDIA, and blot-based assays. Technical challenges such as interpreting glycoRNA topology from whole lysates or organelles, further limit our understanding.

Even though our data identify AGA as a rate-limiting regulator of glycoRNA production, AGA is classically a lysosomal hydrolase that acts late in *N*-glycoprotein degradation, downstream of proteolysis and exoglycosidases such as NEU1.^44^ While purified AGA efficiently cleaves complex, sialylated glycopeptides *ex vitro* and displays broad substrate tolerance, our findings do not yet establish a direct causal mechanism linking lysosomal AGA activity to glycoRNA synthesis *in vitro*. Given the degradative nature of lysosomes, coupling catabolism to RNA glycosylation remains conceptually unexpected and should currently be interpreted as functional association rather than definitive biosynthetic proof.

AGA loss increased global glycoprotein *N*-glycosylation while selectively depleting glycoRNA-associated *N*-glycans, supporting a model in which AGA-released glycans are repurposed for RNA modification. However, we were unable to identify the subcellular compartment or the glycosyltransferase(s) responsible for *N*-glycan attachment to RNA. The identity of a putative “RNA OST” therefore remains unknown. Our preliminary detection of COG complex components in the AGA interactome, together with established retrograde trafficking of lysosomal sorting receptors and vesicular localization of glycoRNA reported by others,^45^ suggests, but does not prove, a trafficking-based coupling mechanism.

While AGA knockout and rescue experiments revealed reciprocal effects on glycoprotein and glycoRNA glycosylation, we cannot exclude broader system-level glycosylation changes caused by chronic gene disruption. However, complementary AGA overexpression analyses can, at least in part, assess dosage dependence and directionality.

Although multiple catalytic and disease-associated AGA mutants were analyzed, detailed structure-function relationships were not explored and remain beyond the scope of this study. Our isotope pulse-chase experiments support delayed glycan incorporation into glycoRNA relative to glycoproteins, consistent with glycan recycling. However, differences in intrinsic biosynthetic efficiency or turnover rates could also contribute. Incorporating AGA-deficient cells and pathway-specific tracers, such as isotope labeled *N*-acetylmannosamine (ManNAc) or sialic acid (Sia), may further resolve these possibilities.

Finally, glycoRNA regulation we found in this work is likely networked rather than linear; for example, NEU1 is unlikely to be essential for glycoRNA biogenesis per se but may modulate glycoRNA glycoform composition by shaping recycled glycan pools. Systematic interrogation of additional lysosomal glycoenzymes will be required to define this regulatory landscape. Together, these limitations highlight both the conceptual novelty of linking lysosomal glycan turnover to RNA glycosylation and the substantial opportunities for future work to define the enzymatic, spatial, and regulatory logic of glycoRNA biogenesis.

In summary, we unravel a mechanistic connection between lysosomal glycoprotein catabolism and RNA glycosylation, identifying AGA as a rate-limiting upstream regulator of glycoRNA biogenesis. These findings redefine lysosomes as active contributors to RNA modification and motivate future studies to uncover additional glycoenzymes in this pathway and to define glycoRNA functions in development, immunity, and disease.

## Methods

### Cell culture, transfection, and stable cell line generation

293T (ATCC) cells, HEK293 (ATCC) cells and all derivative cells, as well as HeLa (ATCC) cells, were cultured in DMEM with high glucose, glutamine and 10% FBS (Biological Industries), 100 units mL^−1^ penicillin and 100 μg mL^−1^ of streptomycin at 37°C in a humidified 5% CO_2_ atmosphere. SLC35A1-KO cells were previously constructed. The GNPTAB-KO (bulk cells), M6PR-KO, SIDT2-KO, SIDT2/1-DKO and AGA-KO cells used in this study were established by the CRISPR/Cas9 system with two different gRNAs, as listed in Table S1.

### RNase treatment and flow cytometry

Cells (5 × 10^5^) were seeded in a six-well plate one day before analysis. The cells were harvest with TrypLE (Thermo Fisher Scientific) for 4 min on ice, divided into three tubes and washed with ice-cold FACS solution (0.5% Bovine serum albumen, 0.1% NaN₃ in 1×PBS). For RNase treatment, cells were resuspended in 100 μL PBS added with 2 μM RNase A/T1 and incubated at 37°C for 30 min, then centrifuged for 2 min at 4°C and 350 × g with 150 μL FACS Buffer. Cells were then stained with 10 μg/mL Siglec-11-Fc for 30 min or 10 μg/mL lectins for 15 min on ice. After incubation, cells were washed twice with FACS Buffer, followed by staining with 10 μg/mL PE-conjugated goat anti-Human IgG antibody or PE-conjugated streptavidin for 30 min or 15 min on ice in the dark. After staining, cells were washed twice with FACS buffer and analyzed using an Accuri C6 (BD, Franklin Lakes, NJ, USA). The resulting data were analyzed with an Accuri C6 and FlowJo software (BD).

### Metabolic chemical reporters and inhibitors

The synthesis of Ac_4_ManNAz, 1,6Pr_2_ManNAz and AMS-ManNAz-P was adapted from a well-documented protocol reported in the literature.^14,15^ Stock solutions of metabolic chemical reporters (MCRs) were made to 100 mM in DMSO (Sigma, USA). In cell experiments all MCRs were used at a final concentration of 100 mM for 48 h. Stock solutions of glycan-biosynthesis inhibitors were all dissolved in DMSO at the following concentrations: 1 mg/mL Brefeldin A (Beyotime, China), 1 mg/mL Swainsonine (Beyotime, China), 5 mg/mL Tunicamycin (Beyotime, China). All inhibitors were used on cells for 48 h at a final concentration of 0.05 μg/mL (Brefeldin A), 2 μg/mL (Swainsonine) and 0.35 μg/mL (Tunicamycin), respectively.

### RNA extraction and purification

RNA extraction and purification were performed according to a previously reported method with minor modifications.^1^ Cells in 10 cm dish were first treated with 1 ml of TRIzol reagent (KeyGEN BioTECH, China) and incubated at 37 °C to denature non-covalent interactions. Phase separation was then conducted by adding 200 μL 100% chloroform, vortexing to mix, and finally spinning at 12,000 rpm for 15 min at 4°C. The upper aqueous phase was carefully transferred to a fresh tube and mixed with equal volume of 100% isopropanol, subsequently incubated for 10 min and centrifuged at 12,000 rpm for 10 min at 4°C to obtain a RNA pellet. The pellet was washed with 1 mL of ice-cold 75 % ethanol, and then redissolved in 40 μL DNase/RNase-free H_2_O. The RNA was then treated on-column with Proteinase K (Promega). Proteinase K is diluted 1:19 in water or DTB and added directly to the column matrix, followed by incubation at 37°C for 45 min. The top of the column was sealed with a cap or parafilm to prevent evaporation. After digestion, the columns were allowed to equilibrate to room temperature for 5 min. The RNA was then eluted by spinning into fresh tubes, followed by a second elution with water. To the combined eluate, 1.5 μg of the mucinase StcE (MCE) is added for every 50 μL of RNA, and the mixture was incubated at 37°C for 30 minutes to allow digestion. The RNA was subsequently cleaned up using a Zymo column. For this cleanup, 2 volumes of RNA Binding Buffer (Zymo Research) were added and vortexed for 10 seconds, followed by the addition of 2 volumes (samples + buffer) of 100% ethanol and vortexing for 10 sec. The mixture was then applied to the column, washed as described above, and eluted twice with water. The final enzymatically digested RNA was quantified using a Nanodrop under the manufacturer’s RNA settings. For small RNA isolation, 2 volumes of adjusted RNA binding buffer (1:1 Zymo RNA Binding Buffer and 100% ethanol) were added to total RNA and vortexed, and the mixture was then applied to a Zymo column. The column flowthrough, which contained the small RNA, was collected, mixed with 2 volumes of 100% ethanol, vortexed, and applied to a fresh Zymo column. The columns were washed twice with 400 μL 80% ethanol, as above, and eluted twice with water.

### Strain-promoted azido-alkyne cycloaddition (SPAAC) reaction to RNA

The extracted RNA labeled by MCRs (50 μg, 36 μL) was mixed with 40 μL ‘‘dye-free’’ Gel Loading Buffer II (df-GLBII, thermoFisher) and 4 μL 10 mM stock of DBCO-PEG4-biotin (Click Chemistry Tools, USA). Samples were conjugated at 55℃ for 10 min in a Thermomixer (Eppendorf) and then purified with as described above.

### Periodate oxidation and aldehyde ligation for glycoRNA labeling

rPAL labeling was performed as previously described.^6^ Briefly, 5 μg of extracted sRNA was lyophilized and redisolved in 28 μL Blocking Buffer (1μL 16 mM mPEG3-Ald (BP-23750, BroadPharm), 15 μL 1 M MgSO_4_ and 12 μL 1 M NH_4_OAc pH5) to block any aldehyde reactive species. The samples were mixed completely by vortexing, and then incubated for 45 minutes at 35℃. The reaction was allowed to cool to room temperature. Then 1 μL 30 mM aldehyde reactive probe (ARP/aminooxy biotin, Cayman Chemicals) was added first, and 2 μL of 7.5 mM NaIO_4_ was also added. The periodate was allowed to perform oxidation for exactly 10 minutes at room temperature in the dark and quenched by adding 3 μL of 22 mM sodium sulfite. The quenching reaction was allowed to proceed for 5 min at 25℃. Next, the reactions were moved back to the 35℃ heat block, and the ligation reaction was allowed to occur for 90 min. The reaction was then cleaned up using a Zymo-I column. 19 mL of water was added in order to bring the reaction volume to 50 mL, and the Zymo protocol was performed as described above.

### RNA gel electrophoresis, blotting, and imaging

Biotin-labeled RNA was resuspended in equal volume of df-GLBII, incubated at 55°C for 10 min and cooled down on ice to denature RNA. Thereafter, 15 µg of RNA was loaded into 1% agarose-formaldehyde denaturing gels, electrophoresed in 1x MOPS buffer (Beyotime, China) at 90 V for 90 min at 4°C. The total RNA in the gel was imaged on a Bio-Rad ChemDoc XRS system (Bio-Rad, USA) after NA-Red staining (Beyotime, China). The imaged gel was then soaked in 20x SSC (Beyotime, China) for two times over 30 min to remove formaldehyde. RNA transfer was adapted from the bottom-up transfer for northern blotting with minor modifications.^46^ Of note, in the transfer process, we chose 10x SSC (Beyotime, China) as the transfer buffer, because its high ionic strength promotes rapid and efficient elution of RNA from the gel while simultaneously facilitating strong electrostatic binding of the negatively charged RNA to the positively charged membrane surface. In the meanwhile, we also opted for positively charged nylon membrane (Cytiva Life Sciences, USA) with higher binding capacity and sensitivity over nitrocellulose membrane, because detecting Biotin-labeled RNA with HRP instead of IR-800 circumvents the strong infrared background intrinsic to nylon membrane. After transfer, RNA was crosslinked to the nylon membrane using UV-light (254 nm, 0.18 J/cm^2^). Nylon membrane was then blocked with Blocking Buffer (Beyotime, China) for 30 min and further incubated with Streptavidin-HRP (ThermoFisher, USA) that was diluted to 1:5000 in Blocking Buffer for 1 h at room temperature. Finally, nylon membrane was washed with Washing Buffer (Beyotime, China) for three times over 20 min and reacted with enhanced chemiluminescence (ECL) reagent (Fdbio Science, China). The bands were detected via a Tanon-5200 chemiluminescence system (Tanon, China).

### Plasmid construction of AGA-Flag and AGA-GFP

All primers, synthetic DNA sequences, and sgRNA sequences used for cloning are listed in Table S1. To knock out target genes with the CRISPR–Cas9 system, single-guide RNAs (sgRNAs) were designed with the E-CRISP website (http://www.e-crisp.org/E-CRISP/), and the targeting DNA fragments were ligated into the BbsI-digested vector pX330-EGFP. To construct pME-AGA-3Flag, the fragments of AGA were amplified from human cDNA library, digested with SalI and NotI, and were cloned into the same site of pME- 3Flag. To construct pME18S-AGA-EGFP, AGA fragment was amplified and assembled into pME18S-EGFP using multi-quick cloning kits.

### Immunofluorescence

HEK293 cells were cultured in glass bottom dishes (Cellvis, CA, USA) to 60% confluence and given fresh culture medium before transfection. The plasmid of pME18S-AGA-EGFP was transfected with the Lipo8000 Transfection Reagent (Beyotime, China) into the cells for protein expression. After 48 hours, the medium was removed and the cells were washed three times with DMEM. Then the cells were stained with Lyso-Tracker Red (50 μM, Beyotime), ER-Tracker Red (1 μM, Beyotime) or Golgi-Tracker Red (0.3 mg/mL, Beyotime) according to manufacturer’s instructions, respectively. Next, the cells were washed with PBS and stained with Hoechst 33342 (10 μg/ml, KeyGEN BioTECH). Finally, the cells were washed with PBS and imaged using an Olympus IXplore SpinSR10 Spinning Disk Confocal Super Resolution Microscope (Olympus Corporation, Japan).

#### The hydrolytic activity of AGA toward the GlcNAc-Asn glycosidic bond in various substrates

For GlcNAc-Asn, 25 μL of 500 μM substrate were incubated with 25 μL of 0.25 ng/μL AGA (SinoBiological, China) in 100 μL Assay Buffer (50 mM Sodium phosphate pH 6.0, 150 mM NaCl, 5 mM DTT) at 37℃ for different lengths of time. followed by centrifugation and filtration to remove protein precipitates. The samples were next analyzed via LC-MS (Agilent 1290 Infinity II system coupled to a G6530 Q-TOF mass spectrometer). Chromatographic separation was performed on an Agilent HC-C18(2) column (4.6 × 250 mm, 5 μm particle size). The mobile phase consisted of (A) 0.1% formic acid in water and (B) 0.1% formic acid in acetonitrile, at flow rate of 1 mL/min with the following gradient program: 95% - 0% A (0-10 min).

For SGP, 50 μL of 500 μM substrate were incubated with 50 μL of 0.25 ng/μL AGA in 200 μL Assay Buffer at 37℃ for 12 h, followed by centrifugation and filtration to remove protein precipitates. The samples were next analyzed via HPLC (Waters 2998 Photodiode Array detector, USA). Chromatographic separation was performed on a Waters XBridge^®^ Peptide BEH C18 column (19 × 150 mm, 5 μm particle size). The mobile phase consisted of (A) 0.1% formic acid in water and (B) 0.1% formic acid in acetonitrile, at flow rate of 7 mL/min with the following gradient program: 95% - 0% A (0-30 min).

For 293 protein or sRNA, protein extraction and sRNA isolation were performed as described above. Then, 2 mg of 293 protein or 5 μg of 293 sRNA were incubated with different amounts of AGA in 200 μL Assay Buffer at 37 ℃ for 12 h, followed by PGC SPE purification to LC-MS/MS analysis as described above.

### CAT-Lyso for in situ lysosomal proteomics

Lysosomal proteomics with LysoCat was performed according to a previously reported method.^19^ 293 cells or HeLa cells at 90% confluence in 10-cm dishes were incubated with or without 5 ml DMEM containing 50 μL 1 mM stock of LysoCat for 30 min at 37 °C. The cells were washed with DMEM three times to remove extra LysoCat, followed by PBS two times to remove phenol red. Then cells were incubated with 5 mL phenol red-free DMEM containing 50 μL 5 mM stock of SF2 probe for 30 min at 37°C. Subsequently, the cells were irradiated by 520 nm green light (the 10-cm dish was covered by an ice box for cooling) for 15 min at r.t. and placed in the dark for 15 min at 37°C. Then the culture medium was discarded and the cells were washed with PBS. After that, the cells were lysed with 1 mL of RIPA strong lysis buffer (Meilunbio, China) containing with 1x protease inhibitor cocktail (TargetMol, China). The sample was transferred into a new 1.5 mL tube and sonicated on ice to generate a clear lysate and the lysate was centrifuged at 12,000 rpm for 20 min at 4°C to remove cell debris. After adjusting each sample to the same protein amount with the BCA assay kit (Pierce, USA), The lysate was next precipitated with methanol-chloroform (MeOH: dissolved protein: CHCl_3_ = 4:2:1, v/v/v), kept at -20°C for at least 1 h to allow for complete precipitation, followed by centrifugation (12,000 rpm, 20 min, 4°C). The sample was then washed three times with cold methanol for subsequent enrichment.

High-capacity streptavidin agarose resin beads (ThermoFisher, USA) were pre-equilibrated by washing three times with PBS. The washed protein was redissolved in 600 μL 2% SDS/PBS, sonicated and heated to 95°C for 10 min to ensure complete solubilization. Each sample was then transferred to a 15-mL centrifuge tube (Corning, USA) and diluted with PBS to a final SDS concentration of 0.2%, followed by incubation with 70 µL streptavidin beads (per 10 mg of protein) at 4°C overnight with gentle shaking. After incubation, the beads were pelleted (1,400x g, 3 min, RT), resuspended in 0.2% SDS/PBS and rotated for 10 min at RT. Each sample was then washed three times with 6 mL 0.2% SDS/PBS and eight times with 6 mL PBS. After the final wash, the beads were pelleted (1,400x g, 3 min, RT) and resuspended in 500 μL PBS to transfer into a 1.5 mL tube (Eppendorf, UK), followed by centrifugation (1,400x g, 3 min, r.t.) to collect the bead pellets. Each bead pellet was then resuspended in 500 μL 6 M urea/PBS, supplemented with 25 μL 200 mM dithiothreitol (DTT), and incubated at 37°C for 30 min with shaking in a ThermoMixer (Eppendorf, UK). Then, 25 μL 400 mM iodoacetamide (IAA) was added and the mixture incubated again at 35°C for 30 min in the dark. After incubation, each sample was diluted with 950 μL PBS and the beads were collected by centrifugation (1,400 g, 3 min, r.t.), with the supernatant removed. Finally, each bead pellet was resuspended in 100 μL of 2 M urea/PBS containing 4 μL trypsin (0.5 μg μL⁻¹) and digested at 37°C for 18 h with shaking. The supernatant was collected. The bead pellet was washed twice with 400 μL PBS, and all supernatants were pooled and dried in a vacuum concentrator.

### Immunoprecipitation

HEK293 cells were cultured in 10-cm dishes to 90% confluence and given fresh culture medium before transfection. The plasmid of pME18S-AGA-Flag was transfected with the Lipo8000 Transfection Reagent (Beyotime, China) into the cells for protein expression. After 48 hours, the culture medium was discarded and the cells were washed with PBS. After that, the cells were lysed with 1 mL of IP lysis buffer (Beyotime, China) containing with 1 mM PMSF (Beyotime, China). The lysate was transferred into a new 1.5 mL tube and centrifuged at 12,000 rpm for 20 min at 4°C to remove cell debris. After adjusting each sample to the same protein amount with the BCA assay kit (Pierce, USA), 2 mg of protein lysates were incubated with 10 μL anti-Flag magnetic beads (ThermoFisher, USA) at 4℃ for 4 hours. After incubation, the immuno-complexes were washed six times with NETN buffer 20 mM Tris-HCl (pH 8.0), 100 mM NaCl, 1 mM EDTA, and 0.5% NP-40). Subsequently, each sample was resuspended in 210 μL Elution Buffer (50 mM Tris-HCl pH 8.0, 10 mM EDTA, 1% SDS) and shaken in a ThermoMixer at 65°C for 30 min. The supernatant was collected. The beads were washed twice with 400 μL PBS, and all supernatants were pooled and dried in a vacuum concentrator.

Subsequently, protein was digested following the S-trap manufacturer’s protocol with minor changes and as we have previously reported.^47^ 23 µL of the lysis buffer (5% SDS, 50 mM triethylammonium bicarbonate (TEAB), pH 8.5) was added to the sample, followed by the addition of 2 µL of 550 mM dithiothreitol (DTT) and incubation at 55 °C for 50 min. Then, 4 µL of 450 mM iodoacetamide (IAA) was added, and the mixture was incubated in the dark for 20 min. After alkylation, the sample was acidified with approximately 4 μL of 27.5% phosphoric acid to achieve a pH below 1.0. The sample was supplemented with 165 μL of binding buffer (100 mM TEAB in 90% methanol), loaded onto an S-Trap column (ProtiFi, USA), and centrifuged at 4,000 × g for 30 s. The column was subsequently washed three times with 150 μL of wash buffer (100 mM TEAB in 90% methanol), centrifuged at 4,000 × g for 1 min, followed by an additional centrifugation to remove residual buffer. For digestion, 2 μg of sequencing-grade trypsin (OMICSOLUTION, China) in 50 mM TEAB buffer was added to the column and incubated at 37°C for overnight. Peptides were sequentially eluted from the S-Trap column by centrifugation (4,000 × g, 1 min per step) using three elution buffers: 40 μL of 50 mM TEAB (Elution 1), followed by 40 μL of 0.1% (v/v) formic acid (FA) (Elution 2), and finally 40 μL of 0.1% FA in 50% (v/v) ACN (Elution 3). The pooled eluates were vacuum-dried using SpeedVac (Labconco).

### MS-based proteomic analysis

Dried peptides were reconstituted in 0.1% FA and centrifuged at 12,000 × g for 15 min to remove insoluble material. The supernatant was transferred to a MS vial and analyzed on a timsTOF Pro 2 mass spectrometer (Bruker Daltonics, Germany) coupled to a nanoElute nanoflow UHPLC system (Bruker Daltonics, Germany) in data-independent acquisition (DIA)-PASEF mode. Peptides were separated on a C18 reversed-phase column (25 cm × 75 μm, 1.9 μm particles; OMICSOLUTION, China) at 40 °C using a 60-min nonlinear gradient at 300 nL/min. Mobile phase A consisted of 0.1% FA in water and phase B consisted of 0.1% FA in ACN, with the gradient ramped from 2% to 22% B over 40 min, increased to 37% B at 50 min, and then raised to 80% B from 55 to 60 min. Ions were analyzed over a range of 100–1700 m/z with trapped ion mobility spectrometry (TIMS) enabled. The TIMS ion mobility range was set from 1/K₀ = 0.60 to 1.60 Vs·cm⁻², with the ramp time set to 100 ms. The CaptiveSpray source was operated at a capillary voltage of 1400 V, and the dry gas flow was set to 3.0 L/min at 180 °C. Ions were transferred with an ion energy of 5.0 eV and fragmented using a collision energy of 10.0 eV at 60 μs. All raw MS files were analyzed with DIA-NN (version 1.9.1).

### 13C glucose pulse-chase and reverse pulse-chase experiment

HEK293 cells were cultured in 10-cm dishes to 50% confluence and given fresh glucose-free DMEM media supplemented with 10% (v/v) FBS and 4.5 g L^−1^ of [UL-^13^C] D-glucose (^13^C glucose, Bide Pharmatech Ltd., China). The cells were washed and collected at 12, 24 and 48 hours, respectively. For ^13^C/^12^C pluse chase experiment, HEK293 cells at 50% confluence were cultured with fresh glucose-free DMEM media supplemented with 10% (v/v) FBS and 4.5 g L^−1^ of ^13^C glucose. After 24 h, the cells were washed with PBS three times and recultured with 4.5 g L^−1^ of ^12^C glucose-contained DMEM media supplemented with 10% (v/v) FBS for different lengths of time. The cell samples were washed and harvested at expected time points. The collected samples were then subjected to proteins extraction and RNA isolation for quantifying the amount of ^13^C labeled monosaccharides in the glycans by LC-MS/MS analysis.

### MS-based glycomic analysis

*N*-glycan release and purification were performed according to a previously reported method.^24^ Brefily, RNA or protein samples were resuspended in 100 μL glycan release buffer (100 mM HEPES, 10 mM DTT, pH 7.5), and denatured at 100℃ for 2 min in a thermomixer (IKA, Germany). After cooling to the room temperature, 4 μL PNGaseF (500 U/μL; NEB) was added to the sample, followed by incubation overnight at 37°C. Following PNGaseF cleavage, the released N-glycans were desalted using a 96-well plate Hypercarb porous graphitized carbon (PGC) solid-phase extraction (SPE) cartridge (Thermo Fisher Scientific, USA). Before desalting, 400 μL 0.1% trifluoroacetic acid (TFA) was added to the sample to adjust the volume. The SPE was preconditioned with 100% acetonitrile (ACN), followed by 80% ACN containing 0.1% TFA, and then equilibrated with 0.1% TFA. The sample was loaded onto the SPE and washed five times with 0.1% TFA. N-glycans were eluted sequentially with 200 μL of 40% ACN in 0.1% TFA and then 200 μL of 80% ACN in 0.1% TFA. The sample was subsequently dried using a CentriVap Centrifugal Concentrators (Labconco, USA).

For MS analysis of released glycans, dried samples were reconstituted in 20 μL of 0.1% (v/v) formic acid (FA) in MS grade water, and 4 μL was injected for LC-MS/MS analysis. Glycan separation was performed on a nanoAcquity UPLC system (Waters) using a Hypercarb porous graphitic carbon column (1 × 100 mm, 3 µm; Thermo Scientific, USA) at 50 µL/min, with mobile phases consisting of A: 0.1% (v/v) formic acid (FA) in water and B: 0.1% (v/v) FA in 80%ACN. The following chromatographic gradient was employed: 0–2 min, 2% B; 2–10 min, 2–12% B; 10–44 min, 12–72% B; 44–45 min, 72–100% B; 45–50 min, 100% B; 50–51 min, 100–0% B; and 51–60 min, 0% B. MS analysis was performed in positive SWATH mode using a ZenoTOF 7600 system (SCIEX) with the Zeno trap on. Ion source gas1, gas2, and curtain gas were at 20, 45, and 35 psi, respectively. MS1 spectra were acquired across m/z 500–2000 with a 200 ms accumulation time. Glycan fragmentation employed dynamic nitrogen gas collision-induced dissociation in 48 SWATH windows. Product ions were monitored over an *m/z* range of 140–1800, with an accumulation time of 15 ms SWATH window. Glycan identification and quantitation were conducted using GlycanDIA Finder and Skyline (V25.1). Integrated peak areas were extracted from high-resolution LC-MS/MS data and used as quantitative metrics of glycan abundance. Relative abundances of individual glycan compositions were calculated by normalizing each peak area to the total peak area of all identified glycans.

### RNA acp^3^U modification analysis

RNA modification analysis was performed as previously described.^26,48^ Brefily, 1 μg purified small RNAs were mixed with 20 μL of digestion buffer (1 mM ZnCl₂, 1 mM MgCl₂, 30 mM sodium acetate, pH 7.2) containing 5 mU/μL nuclease P1 (Sigma-Aldrich), 5 mU/μL shrimp alkaline phosphatase (New England Biolabs), 500 μU/μL phosphodiesterase I (Sigma-Aldrich), and 6.25 μU/μL phosphodiesterase II (Worthington Biochemical), and the reaction was carried out overnight at room temperature. The digested nucleoside was purified using Hypercarb PGC SPE 96-well plates (Thermo Fisher Scientific) performed as described above.

For MS analysis of digested RNA, 2 μL of each sample was injected and separated using a nanoAcquity UPLC system (Waters) equipped with a Hypercarb PGC column (1 × 100 mm, 3 μm; Thermo Scientific) at a constant flow rate of 50 μL/min. The mobile phases consisted of (A) 0.1% FA in water and (B) 0.1% FA in ACN with the following gradient: 0–2 min, 0% B; 2–20 min, 0–16% B; 20–40 min, 16–72% B; 40–42 min, 72–100% B; 42–52 min, 100% B; 52–54 min, 100– 0% B; 54–65 min, 0% B for column re-equilibration. Nucleosides were ionized using a ZenoTOF 7600 mass spectrometer (SCIEX) in positive electrospray mode with the Zeno trap on. The source parameters were set as follows: curtain gas at 35 psi, ion source gas 1 at 40 psi, ion source gas 2 at 60 psi, and ion transfer tube temperature at 200°C. acp^3^U and U were monitored using a targeted method at 346.12 and 245.08 m/z, respectively, and modifications were identified and quantified using Skyline (version 25.1).^49^

### Lysosome isolation

Lysosomes were isolated using the Minute^TM^ Lysosome Isolation Kit for mammalian cells/tissues (LY-034, invent): Cells were cultured in 10-cm dishes to 90% confluence and washed three times with ice-cold PBS. In the third PBS wash, cells were scraped off the plate and spun down at 500x g for 5 min at 4℃. Cell pellets were resuspended in 500 μL of Buffer A per 25,000,000 cells. After incubating the suspension on ice for 10 min, cell suspension was vigorously vortexed for 20 seconds and immediately transferred into the filter cartridge. The cartridge was capped, inverted several times, and centrifuged at 16 000 × g for 30 s. The flow-through was resuspended and re-passed through the same filter to increase yield. After discarding the filter, the retentate was resuspended by vigorous vortexing for 10 s, and the tube was centrifuged at 2 000 × g for 3 min to pellet nuclei and debris.The supernatant was transferred to a fresh 1.5-mL microfuge tube and spun at 11 000 × g, 4 °C for 15 min. About 400 µL of the resulting supernatant was carefully removed, placed in a new tube, and centrifuged at 16 000 × g, 4 °C for 30 min, after which the supernatant was completely aspirated. The pellet was resuspended in 200 µl of cold buffer A by pipetting up and down 80 times and vigorous vortexing for 20 s, centrifuged at 2 000 × g for 4 min, and the supernatant was transferred to a clean 1.5-mL tube again. 100 µL of buffer B was added (2:1 ratio). The mixture was briefly vortexed, incubated on ice for 30 min, and centrifuged at 11, 000 × g for 10 min. All supernatant was removed, and a short re-spin at 11 000 × g was performed to eliminate residual buffer, obtaining a pellet as lysosome.

### Methylated RNA Immunoprecipitation Sequencing (MeRIP-seq)

MeRIP-seq and data analysis were done by Guangzhou Epibiotek Co., Ltd. The isolated lysosomes were subsequently used for RNA extraction as described above. The concentration of total RNA was measured by Qubit RNA HS assay kit (Invitrogen, Q32852). The poly(A)+ RNA from total RNA was isolated using oligo(dT) coupled to magnetic beads. 3 μg poly(A) + RNA was fragmented into 100-200 nt RNA fragments using 10X RNA Fragmentation Buffer (100 mM Tris-HCl, 100 mM ZnCl_2_ in nuclease-free H_2_O). The reaction was stopped by adding 10X EDTA (0.5 M EDTA). Methylated RNA immunoprecipitation was performed using Epi^TM^ m6A immunoprecipitation kit (Epibiotek, R1802). Briefly, the fragmented RNA was incubated with anti-m6A monoclonal antibody (abcam, ab208577) for 3 h at 4℃ and then with protein A/G magnetic beads (Invitrogen, 8880210002D/10004D) at 4℃ for an additional 2 h to obtain immunoprecipitated RNA fragments.

The m6A-enriched RNA was purified using TRIzol^TM^ Reagent (Invitrogen, 15596018). The library was prepared by smart-seq method. Both the input samples without IP and the m6A IP samples were subjected to 150-bp, paired-end sequencing on Next-Generation Sequencing (NGS) platform.

### Acetylated RNA Immunoprecipitation Sequencing (acRIP-seq)

AcRIP-seq and data analysis were done by Guangzhou Epibiotek Co., Ltd. Lysosomal total RNA was frag-mented into 100–200 nt RNA fragments using 10X RNA FragmentationBuffer (100 mm Tris-HCl, 100 mm ZnCl_2_ in nuclease-free H_2_O). The reaction was stopped by adding 10X EDTA (0.5 M EDTA). To obtain immuno-precipitated RNA fragments, fragmented RNA was incubated for 3 h at 4°Cwith anti-ac4C monoclonal antibody and then for 2 h at 4°C with protein A/G magnetic beads (Cat# 8880210002D/10004D, Invitrogen) according to the EpiTM ac4C immunoprecipitation kit (Epibiotek, R1815). The library was prepared using the smart-seq method. Both the input samples without IP and the ac4C IP samples were subjected to 150-bp, paired-end sequencing on Next-Generation Sequencing (NGS) platform.

## Acknowledgments

We thank Prof. Dr. Long Liu, Dean of the School of Biotechnology, Jiangnan University, for encouraging and providing financial support for young researchers. We are grateful to Heideki Nakanishi, Yicheng Wang, and Weiwei Ren for insightful discussions. We thank Meiru Lu, Mingyu Li, Yao Liu, and Lekang Wang for assistance with the expression and purification of Siglec11-Fc, and Linpei Zhang for assistance with cell sorting and FACS analysis. We also thank the MS platforms of the Core Facility of Shanghai Medical College, Fudan University, for their support in glycomic and proteomic analyses. This manuscript was shown to Prof. Dr. Xing Chen, faculty of Department of Chemical Biology, College of Chemistry and Molecular Engineering in Peking University, and Prof. Dr. Ryan A. Flynn, faculty of Stem Cell Program and Division of Hematology/Oncology, Boston Children’s Hospital, Department of Stem Cell and Regenerative Biology and Harvard Stem Cell Institute, Harvard University, and received with insightful and ebullient feedback.

We gratefully acknowledge support from the National Natural Science Foundation of China (22477055, 92478107 and 22507114), the China Postdoctoral Science Foundation (2024T170350), the Natural Science Foundation of Jiangsu Province (BK20232020), the National Key Research and Development Program of China (2022YFA1505600), the STI2030-Major Projects (2022ZD0211804), the Greater Bay Area Institute of Precision Medicine (I0036(A)), the Young Talent Program, SCF of Jiangsu Province and the Open Research Fund of Beijing National Laboratory for Molecular Sciences.

## Author contributions

Y.L. conceived the project. R.X., Y.L., Y.X. and M.F. supervised the project and obtained funding. W.C. conducted the glycoRNA labeling experiments. Z.Y., Y. M., S.Z., C.Y. and X. Z. constructed gene-knockout cell lines. L.Y., T.L., Y.Z., T.L., J.J., T.W., and Y.D. performed glycanDIA experiments and analyzed the data. Y. X., R.X and Y.L. wrote the manuscript. All authors discussed the results and revised the manuscript.

## Data availability

Data are available in the main Figures, Extended Data Figures, and Supplementary Figures. The raw mass spectrometry data generated in this study have been deposited in the Mass Spectrometry Interactive Virtual Environment (MassIVE) database under accession code MSV000101822 and MSV000101786. Any additional requests for information can be sent to the lead contact.

## Extended Data Figure Legends

**Figure S1.**
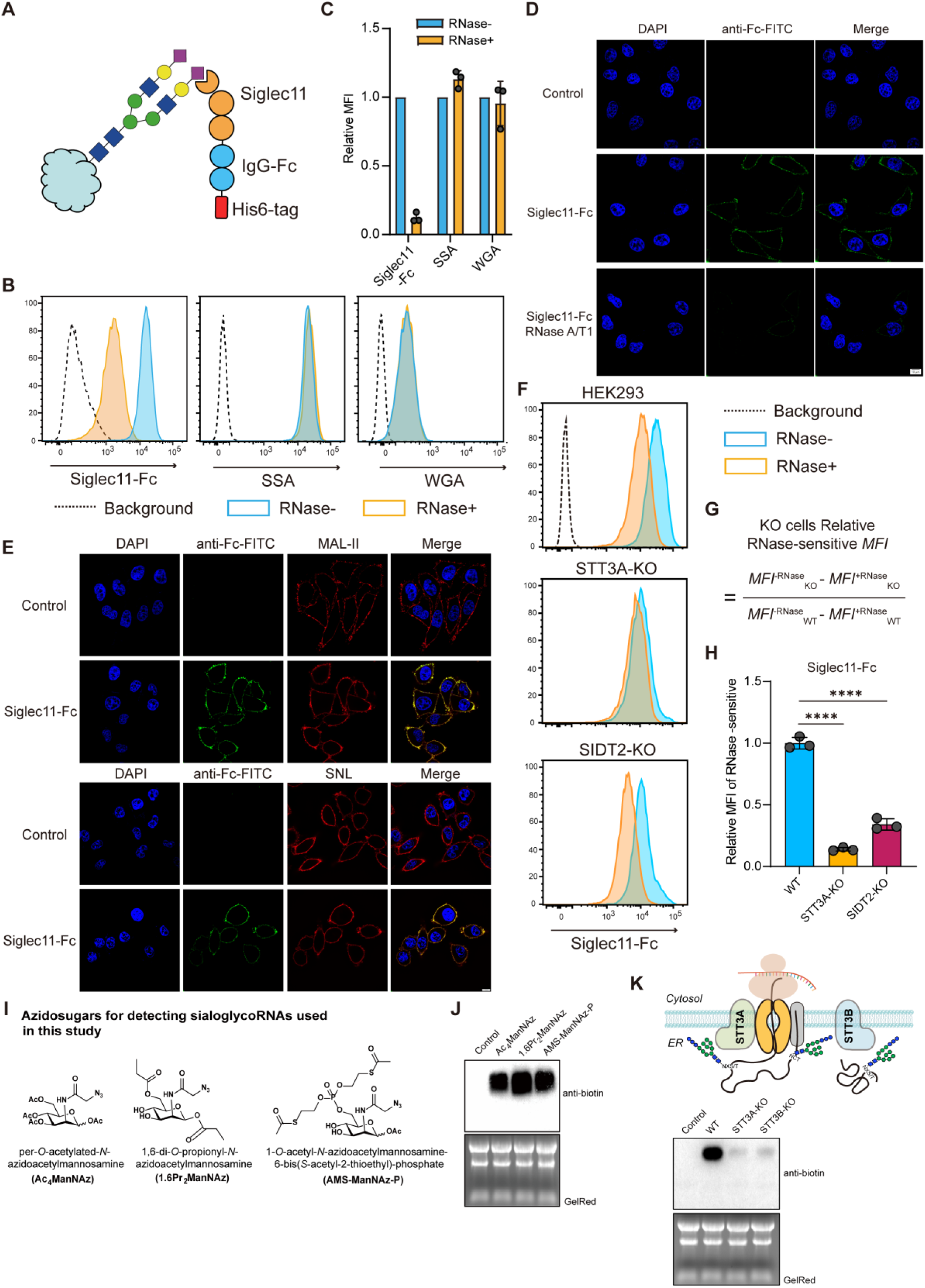
Screening assay and identification of lysosomal trafficking proteins in sialoglycoRNA production. **(A)** Cartoon of recombinant Siglec11-Fc expressed in *E. coli*. **(B)** Flow cytometry of HEK293 cells treated with or without RNase and stained with Siglec11-Fc, SSA, or WGA. **(C)** Quantification of flow cytometry data in (B), WT MFI was set to 1 (mean ± SD, *n* = 3). **(D)** Representative confocal images of RNase-treated or untreated cells stained with Siglec11-Fc (FITC channel, λex = 488 nm). Scale bar, 10 μm. **(E)** Confocal images of Siglec11-Fc co-stained with SNL or MAL-II to indicate sialoglycoproteins. Scale bar, 10 μm. **(F)** Flow cytometry histograms of STT3A-KO or SIDT2-KO cells stained with Siglec11-Fc. **(G)** Algorithm-based analysis of Siglec11-Fc staining differences in KO strains after RNase treatment. **(H)** Quantification of flow cytometry data in (F). WT MFI was set to 1 (mean ± SD, *n* = 3). **(I, J)** MCRs and RNA blotting of HEK293 RNA labeled with Ac_4_ManNAz, 1,6-Pr_2_ManNAz, or AMS-ManNAz-P. Statistical significance was determined by two-tailed Student’s *t*-test: \*\*\*\**P* ≤ 0.0001.

**Figure S2.**
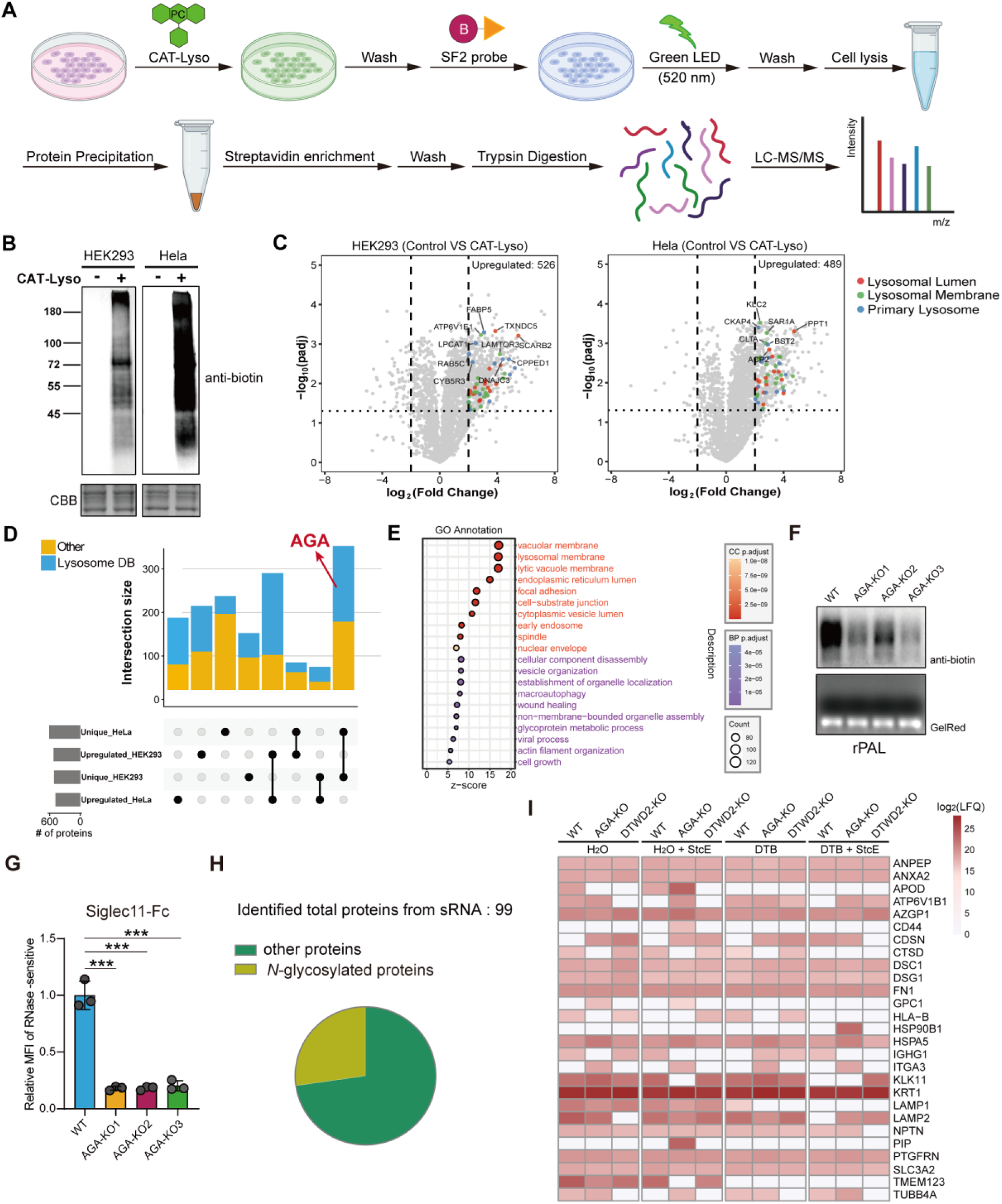
CAT-Lyso profiling of the lysosomal proteome in HEK293 and HeLa cells. **(A)** Workflow of CAT-Lyso labeling coupled with LC-MS/MS for proteomic analysis. **(B)** Western blot of CAT-Lyso biotin labeling in HEK293 and HeLa cells; protein loading normalized by Coomassie Brilliant Blue (CBB) staining. **(C)** Volcano plots showing proteomic changes in the lysosome from three independent replicates; proteins with *P* < 0.05 and fold change > 1.5 (red) were considered upregulated. **(D)** Number and overlap of proteins upregulated or uniquely identified in HEK293 and HeLa cells. **(E)** Gene Ontology (GO) analysis of proteins identified in HEK293 and HeLa cells by CAT-Lyso. **(F)** rPAL blotting of small RNA samples extracted from HEK 293 WT or triplicate biological AGA-KO clones. **(G)** Quantification of flow cytometry data of HEK 293 WT or triplicate biological AGA-knockout clones stained with Siglec11-Fc. WT MFI was set to 1 (mean ± SD, *n* = 3). **(H)** Identified protein quantity and glycoprotein proportion from sRNA extracted from HEK 293 WT, DTWD2-KO or AGA-KO clone treated with 100 mM Ac_4_ManNAz for 48h, which were subjected to the conventional nondenaturing proteinase K treatment or to the modified proteinase K treatment in DTB, and then further treated with mucinase StcE protease. **(I) (I)** Heatmap of identified glycoprotein under different RNA purification conditions from (H). Statistical significance was determined by two-tailed Student’s *t*-test: \*\*\**P* ≤ 0.001.

**Figure S3.**
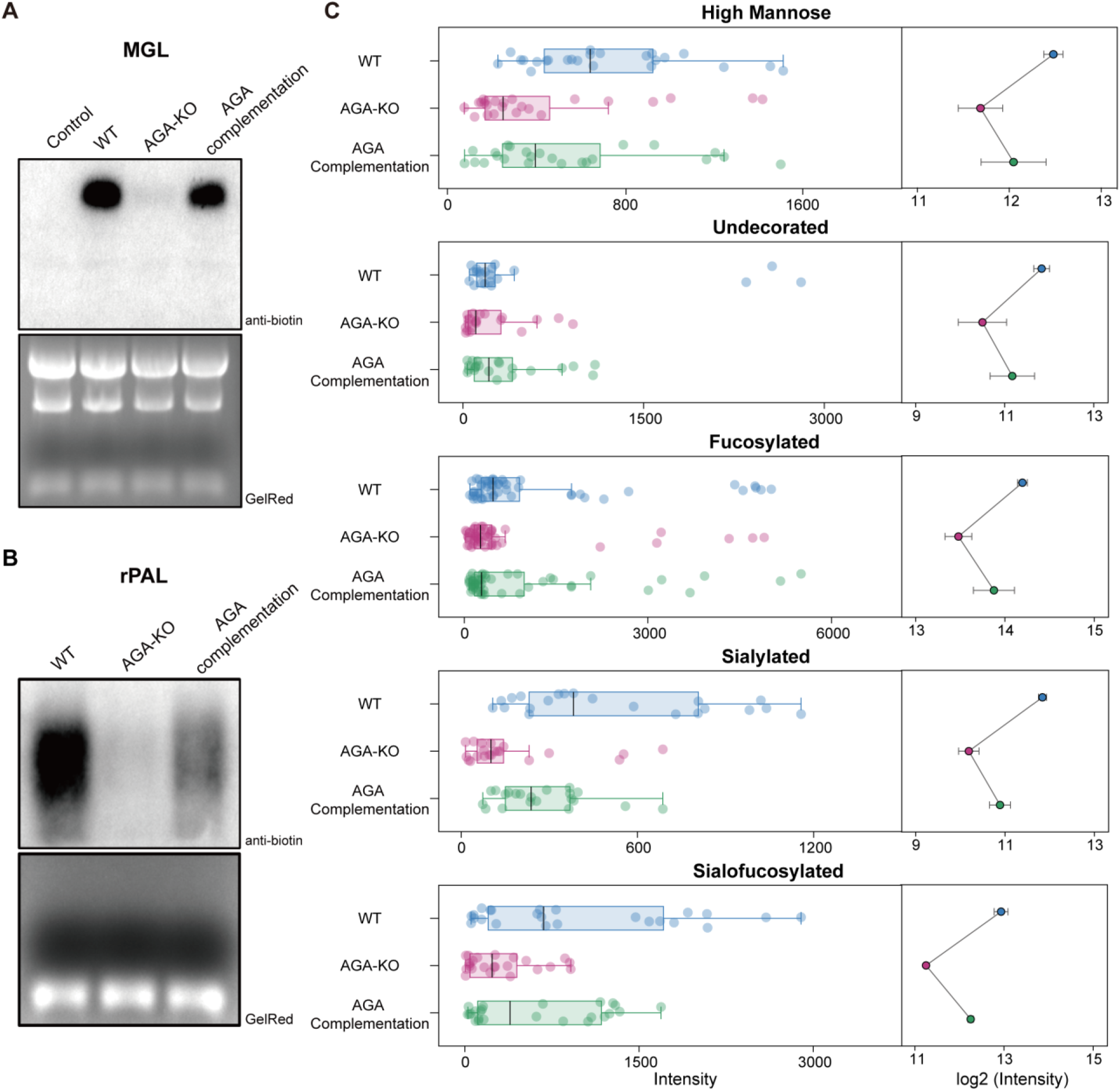
Effects of AGA Complementation and Site-Specific Mutation. **(A)** RNA blotting of RNA from HEK293 WT, AGA-KO, or AGA-complemented cells treated with 100 mM Ac_4_ManNAz for 48 h. **(B)** rPAL blotting of small RNAs from the corresponding cells. **(C)** Relative abundance of five glycan subtypes in glycoRNAs from HEK293 WT, AGA-KO, or AGA-complemented cells

**Figure S4.**
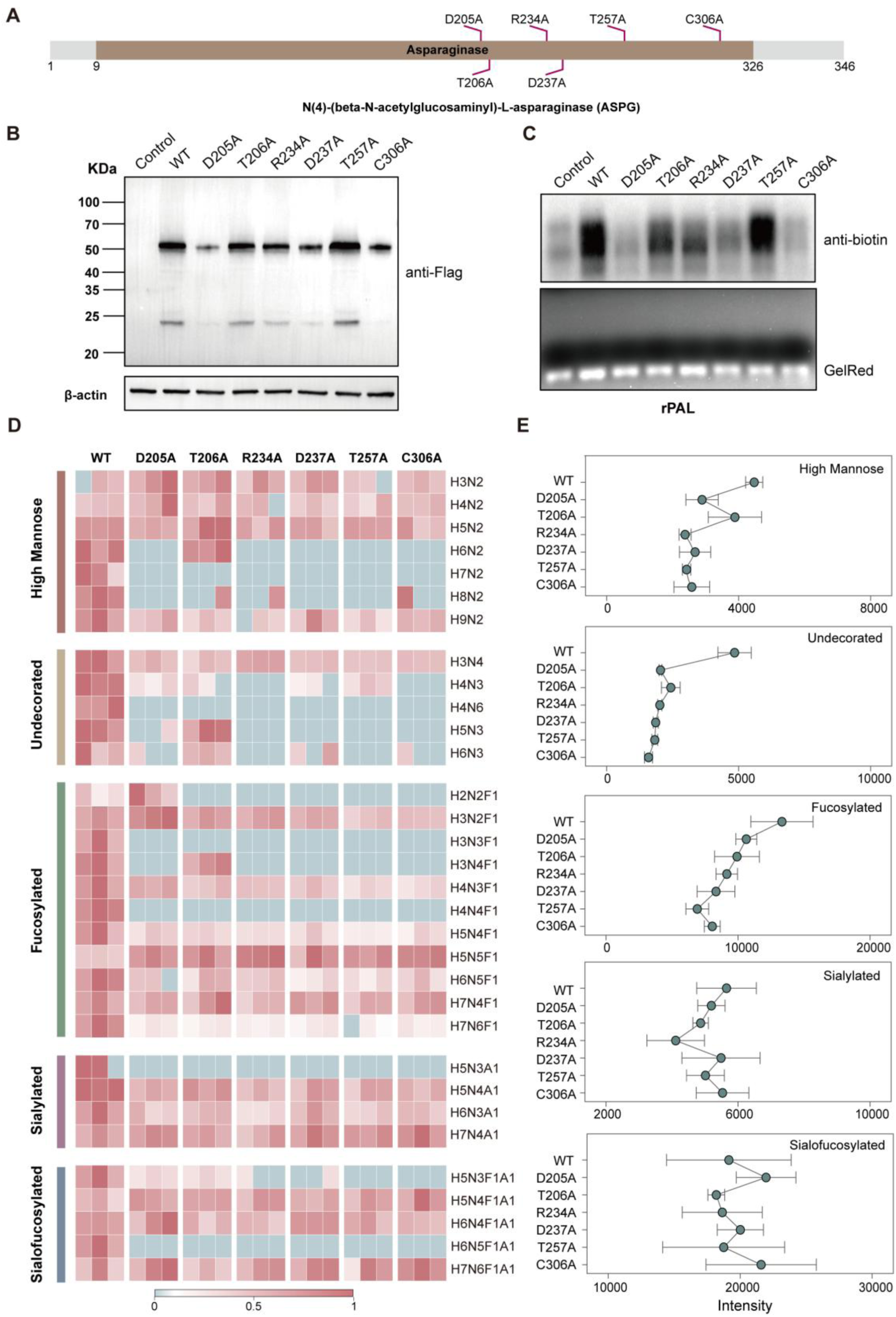
Effects of AGA Site-Specific Mutation. **(A)** Illustration of AGA domain and the selected mutation sites. **(B)** Western blot of HEK293 cells transfected with Flag-tagged AGA plasmids carrying site-specific mutations. **(C)** rPAL blotting of small RNAs from cells expressing mutant AGA plasmids. **(D)** Heatmap of total RNA *N*-glycans identified from cells expressing mutant AGA. **(E)** Relative abundance of five glycan subtypes in glycoRNAs from cells expressing mutant AGA.

**Figure S5.**
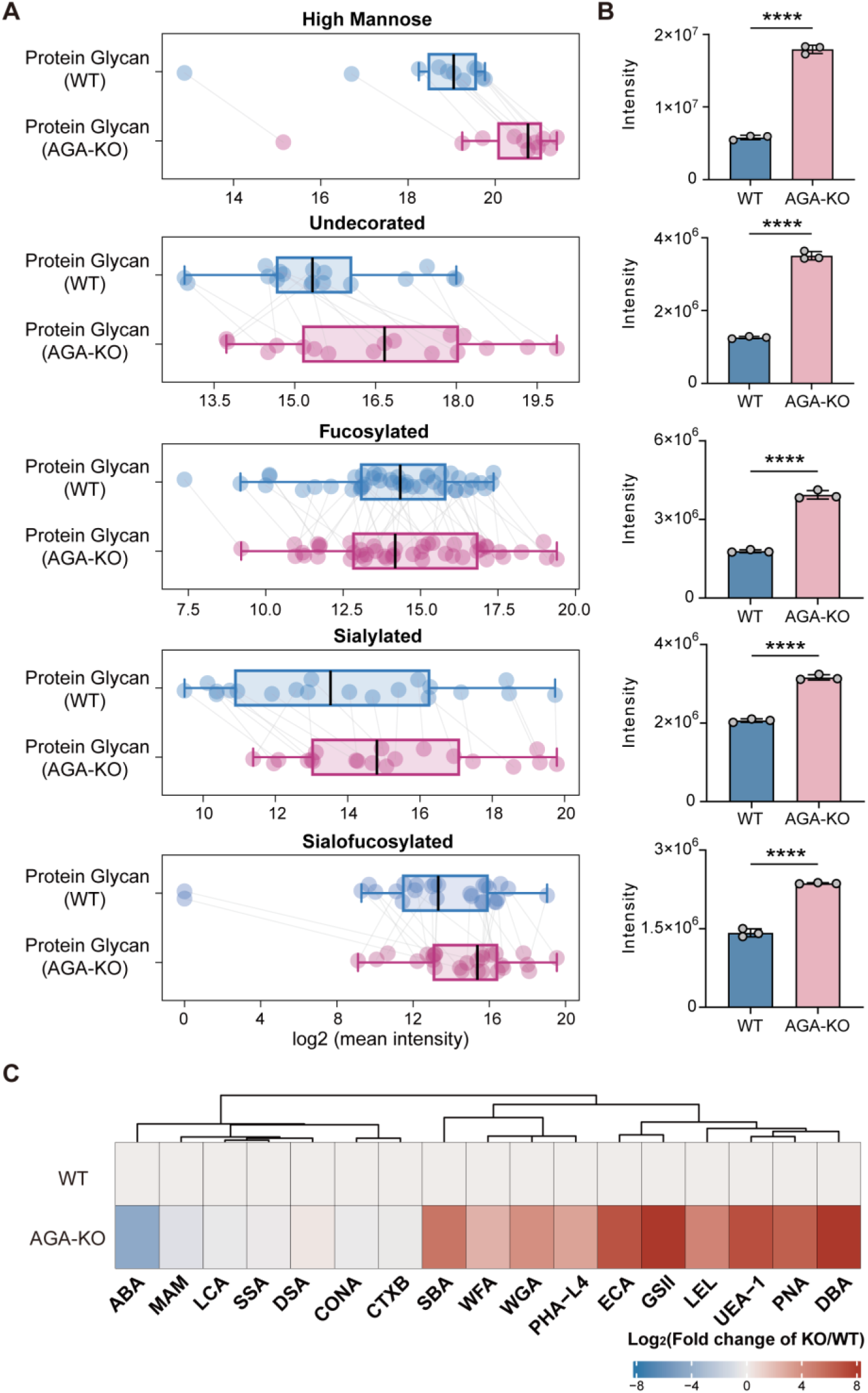
AGA regulates protein *N*-glycosylation in HEK293 cells. **(A, B)** Relative abundance of five glycan subtypes in protein *N*-glycans from WT or AGA-KO cells, showing distinct profiles. **(C)** Heatmap of lectin fluorescence signals from WT or AGA-KO cells; data represent mean ± SEM of three independent experiments. Statistical significance: \*\*\*\**P* ≤ 0.0001.

**Figure S6.**
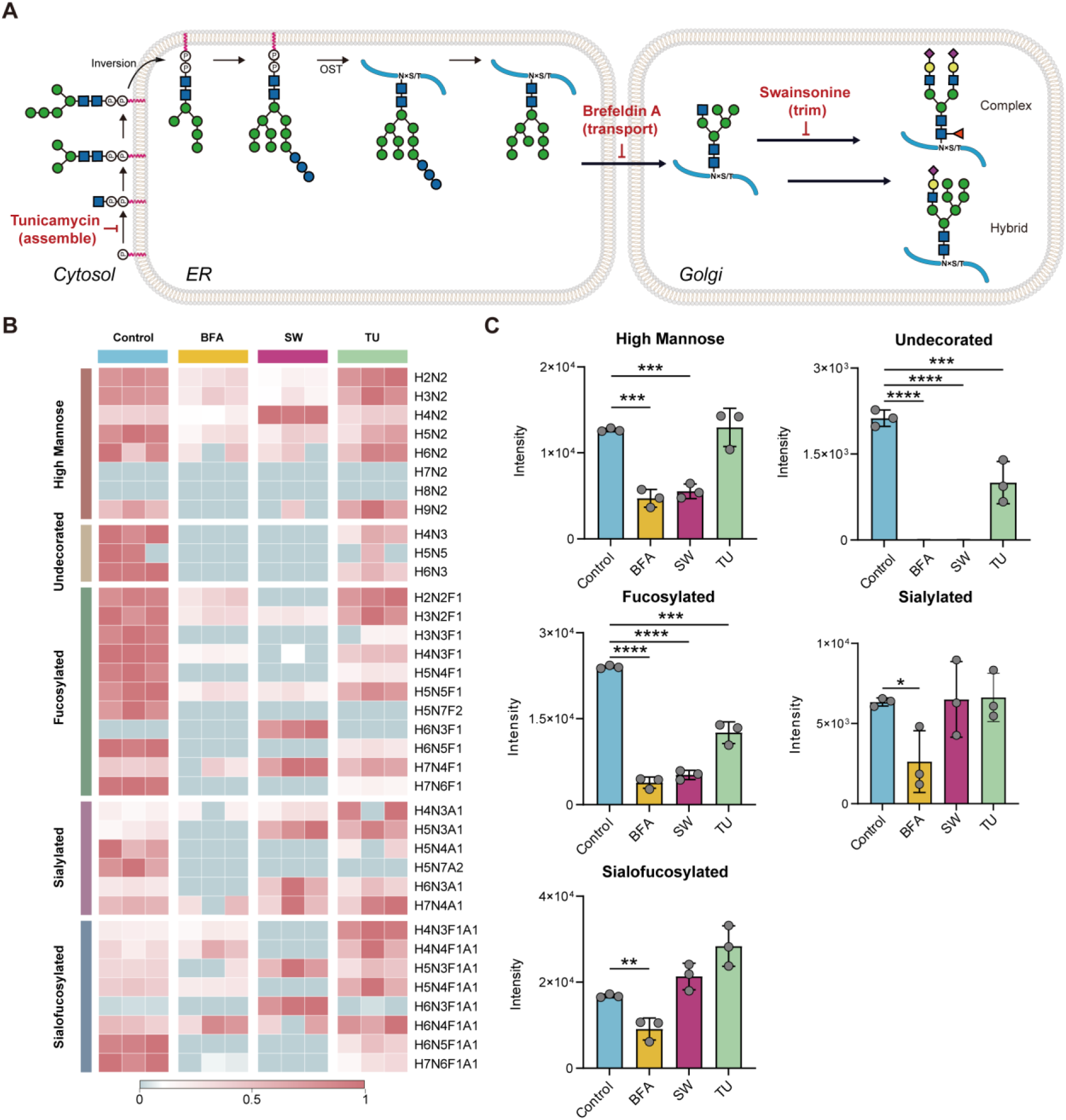
Effects of STT3A/B and Golgi inhibitors on RNA *N*-glycosylation. **(A)** An overview of major steps in the ER- and Golgi-localized steps of N-glycan biosynthesis. Dolichol-linked precursor glycans are transferred to ER-lumen-facing peptides (X) by OST. These glycans are sequentially modified by enzymes that first trim and then elaborate the glycan tree. Specific chemical inhibitors such as Tunicamycin (inhibiting GlcNAc-1-P transferase), Brefeldin A (inhibiting vesicle trafficking/blocking ER-to-Golgi transport) and Swainsonine (inhibiting a-mannosidase II) are employed to interrogate the importance of each of these enzymes on the expression of glycoRNA. **(B, C)** HEK293 WT cells treated with brefeldin A (BFA), swainsonine (SW), or tunicamycin (TU) for 48 h were similarly analyzed, showing altered glycan profiles and subtype distributions. Data represent mean ± SEM of three independent experiments; statistical significance: ns, not significant; \**P* < 0.05; \*\**P* ≤ 0.01; \*\*\**P* ≤ 0.001; \*\*\*\**P* ≤ 0.0001.

